# Resolving the Abstract–Concrete Paradox in the Angular Gyrus: A Multimethod Investigation

**DOI:** 10.64898/2026.02.15.706008

**Authors:** Rocco Chiou, Francesca M. Branzi, Elizabeth Jefferies

## Abstract

Despite its well-established role in memory-guided cognition, whether and how the angular gyrus (AG) contributes to semantic processing remains unresolved. Connectomic work links the AG to abstract mentation, yet neuroimaging studies paradoxically show greater AG engagement for concrete than abstract semantics. To address this discrepancy, we conducted a multimethod investigation integrating neurostimulation, neuroimaging, and experience sampling across five studies (127 human participants; 53 males; 74 females). Using the contrast between concrete and abstract semantics as a diagnostic test-case, we show that this apparent contradiction reflects multiple interacting neurocognitive factors shaping the AG’s functional repertoire. In Study 1, causal disruption of the AG disproportionately impaired abstract semantics and temporary retention of task-relevant information, demonstrating its role in abstract and memory-guided cognition. In Study 2, the apparent concreteness effect was abolished after controlling for performance speed/accuracy, indicating that AG activity is more strongly associated with low-demand mental states than semantic concreteness. In Study 3, the AG showed enhanced functional coupling with regions in the semantic network for abstract words, suggesting sustained integration with the semantic system despite reduced activation. In Study 4, experience sampling showed that concrete words are associated with experiences of mental imagery and automaticity, providing a phenomenological account of heightened AG engagement under low-demand states. Finally, Study 5 clarified how baseline choices influence the interpretation and polarity of AG activity. Together, these findings show that AG engagement is best understood along a continuum of memory-guided cognition, clarifying when and why this region supports abstract versus concrete semantics across situations.

**Significance:** The contribution of the AG to semantic processing has long been debated, both regarding whether it is reliably engaged and how such engagement should be interpreted. By integrating evidence from multiple independent datasets using complementary methodologies, the present investigation moves beyond single-process accounts to identify multiple contributors that jointly define AG function. These factors align with contemporary connectomic research that situates the AG at the transmodal apex of a principal cortical gradient. Under this brain-connectomic framework, AG involvement is amplified as representations become abstract, information is buffered, operation becomes automatic, imagery is simulated, reliance on perception is reduced, shifting from perception-guided mode to memory-guided cognition. Together, these findings provide a principled framework for interpreting the elusive patterns of AG functionality.

---

The angular gyrus (AG) is a parietal region located at the intersection of large-scale brain networks (Seghier, 2013; Humphreys *et al*., 2022), occupying the transmodal apex of a connectomic gradient (Margulies *et al*., 2016). Due to its locus in an intersection position, the AG is thought to support internally oriented cognition, including episodic memory and semantic knowledge (Smallwood *et al*., 2021a). Converging neuroimaging and neurostimulation evidence shows that AG activity tracks phenomenological richness/vividness during episodic recollection, while AG disruption impairs episodic recollection (Yazar *et al*., 2017; Bonnici *et al*., 2018; Rugg and King, 2018; Tibon *et al*., 2019; Kwon *et al*., 2022). Beyond episodic memory, the AG is also engaged by short-term retention of task-relevant information (Lanzoni *et al*., 2019; Murphy *et al*., 2019). Based on these findings, Humphreys et al. (2021) proposed that the AG supports cognition by transiently buffering task-relevant information for online processing.

Despite its established role in memory-guided cognition, AG contribution to semantic processing remains a contentious issue. Although the AG is engaged during sentence comprehension and combinatorial semantics across multiple words (Price *et al*., 2015; Branzi *et al*., 2020), its role in single-word meaning remains debated. Rather than reflecting a unitary function, AG engagement during semantic tasks appears to be shaped by multiple interacting neurocognitive factors that may amplify or counteract each other. Here, we focus on five candidate contributors to AG activation: representational abstractness, information buffering, imagery, automaticity, and choice of baseline. We use the contrast of ‘concrete *vs.* abstract semantics’ as a test-case because connectomic evidence predicts greater AG engagement for abstract cognition, whereas neuroimaging studies often report stronger AG activation for concrete than abstract words. We therefore seek to reconcile this discrepancy by identifying the factors that enhance or dampen AG activity.

## Representational abstractness

Evidence from connectomic research has demonstrated that the AG reliably favours abstract cognition over concrete/tangible perception (Margulies *et al*., 2016). However, neuroimaging studies often report greater AG activation for concrete than abstract words (Hoffman and Bair, 2025). This contradiction is further complicated by multivoxel decoding results that abstract semantics is reliably encoded in the AG, even when univariate activation is weaker (Kaiser *et al*., 2022). This literature indicates an inconclusive pattern regarding whether the AG preferentially contributes to concrete or abstract semantic processing.

## Information buffering

The AG has been proposed to support the retention of information in short-term memory buffer (Humphreys *et al*., 2021). While buffering demand is most pronounced for episodic memory, similar demands arise during semantic tasks when the meaning of multiple words must be combined during reading. This predicts that, if we disrupt the AG with neurostimulation, performance would be more affected when tasks put a high demand on mnemonic retention than situations that rely minimally on buffering.

## Automaticity

Evidence suggests that AG activity scales with proficiency, with greater AG activation observed when performance is facilitated by familiarity or practice (Vatansever et al., 2017). Because concrete words are often processed more easily than abstract words, greater AG responses to concrete semantics may reflect greater automaticity rather than semantic concreteness itself. Disentangling semantic effects from performance-related modulation is therefore critical and is examined in the present study.

## Mental imagery

Concrete concepts (e.g., peacock) are reliably judged to be more imaginable than abstract concepts (e.g., contingency) (Altarriba *et al*., 1999). Given the findings that the AG is one of the areas frequently engaged by imagery (Summerfield *et al*., 2010; Boccia *et al*., 2015), we test whether imagery-related experiences are more likely to arise during the comprehension of concrete than abstract words, providing a phenomenological account of their differential AG engagement.

## Choice of baseline

AG activity is highly sensitive to baseline selection (Humphreys and Tibon, 2023; Fernandino and Binder, 2024). Because passive rest is a mental state potentially semantically abundant, semantic tasks contrasted against rest often appear as *negative deactivation*. In contrast, comparisons against tasks devoid of semantic content typically yield *positive activation*. Moreover, some semantic tasks engage the AG even relative to rest, suggesting that they involve distinctive features capable of driving AG activity to exceed semantically rich resting states. Thus, we also assess how baseline choice might lead to different interpretations of the same result.

Here, we integrate five complementary datasets to characterise how these dimensions jointly shape AG contributions to semantic cognition. Using the contrast between concrete and abstract semantics as a test-case, we test five hypotheses: (*i*) Disrupting the AG with transcranial magnetic stimulation (TMS) would disproportionately impair abstract semantics and tasks requiring online buffering; (*ii*) The apparent concreteness effect in AG activity would substantially diminish after accounting for cognitive effort; (*iii*) Despite ostensibly lower activation, the AG remains connected with semantic-related regions when processing abstract concepts; (*iv*) Processing concrete semantics is associated with imagery-related thoughts and cognitive automaticity; (*v*) the direction and magnitude of AG engagement varies with different choices of baselines.

## Materials and Methods

The present investigation comprises five studies. In Study 1, we used TMS to causally test whether three neurocognitive factors known to modulate AG activity (concreteness, memory buffering, and multimodality) might interact with TMS during a semantic task. In Study 2, with the fMRI data from Chiou et al. (2025), we assessed whether the concreteness effect in AG activity might be an epiphenomenon of task difficulty. In Study 3, with the fMRI data from Hoffman et al. (2015), psychophysiological interaction (PPI) was used to test whether the AG remains integrated with semantic regions during the processing of abstract words. In Study 4, we used MDES (multidimensional experience sampling, a behavioural method for probing thought patterns; Smallwood et al., 2021b) to characterise differences in subjective experiences during the processing of concrete and abstract words, providing phenomenological insight into which facets of thoughts might relate to heightened AG activity for concrete semantics. Finally, in Study 5, we re-analysed multiple datasets, including data from Chiou et al. (2020), to examine how AG engagement might be modulated by baseline choice and task paradigm.

### Participants

#### Study 1

Twenty-four volunteers (11 male, age: 23 ± 2 years) gave informed consent before taking part. All reported right-handedness and English as mother tongue. All participants had normal/corrected-to-normal vision, completed the safety screening for TMS/MRI before the study, and reported no history of neurological disease/injury. This study was reviewed and approved by the institution ethics committee at University of Surrey.

#### Study 2

Twenty-five volunteers (10 male, age: 28 ± 6 years) gave informed consent before their fMRI scans. Participant details of Study 2 are reported in Chiou et al. (2025).

#### Study 3

Eighteen volunteers (10 male, age: 20 – 39 years) gave informed consent before their fMRI scans. Participant details of Study 3 are reported in Hoffman et al. (2015).

#### Study 4

Thirty-six volunteers (16 male, age: 26 ± 5 years) gave informed consent before the MDES testing. All reported right-handedness and spoke English as native language. This study was approved by the ethics review board at University of Surrey.

#### Study 5

Twenty-four participants (6 male, age: 25 ± 7 years) gave informed consent before their fMRI scans. Participant details of Study 5 are reported in Chiou et al. (2020).

### Study 1

Study 1 entailed three experimental sessions at least two days apart. In Session 1, we acquired a high-resolution T1-weighted anatomical image for each participant using a Siemens Trio Scanner. In-plane resolution was 0.8 × 0.8 × 0.8 mm. In Session 2 and 3, we conducted behavioural testing combined with TMS applied to either the AG or the control site vertex (with order counterbalanced across individuals). Visual and auditory stimuli were presented using MATLAB with Psychtoolbox (Brainard, 1997; Pelli, 1997) on a rectangle screen (60.9 × 58.4 cm; 75 Hz refresh rate; 1024 × 768 resolution) and two DELL speakers located next to the screen. Viewing distance was approximately 57 cm from the screen. TMS was applied via a Magstim Super Rapid^2^ system equipped with a figure-of-eight D70^2^ coil. Coil localisation was guided with Brainsight 2 (Rogue Research Inc.), a frameless stereotaxic neuronavigation system that enables a standardised co-registration procedure (e.g., Chiou and Lambon Ralph, 2016b, a; Chiou and Ralph, 2018; Branzi *et al*., 2021) such that each participant’s native T1-weighted structural scan was co-registered with their scalp while the position of TMS coil was constantly adjusted to achieve precise targeting. For AG stimulation, the target was defined using a coordinate from the meta-analysis of Humphreys and Lambon Ralph (2014): *X* =-48, *Y* =-64, *Z* = 34 in the MNI space (Fig. 1A). This site, located in the middle AG near the border of two major subregions, PGa and PGp, of the inferior parietal lobule (Seghier, 2013; Caspers and Zilles, 2018), is a convergence locus identified across many cognitive tasks, including semantic processing, episodic retrieval, and sentence comprehension. This coordinate was chosen due to its intersectional position among multiple networks, wide associations with various domains, and the fact that it has been used to localise regions of interest (ROI) in several fMRI studies (e.g., Humphreys *et al*., 2024). For each individual, this AG coordinate in the MNI space was converted into the corresponding coordinate in each one’s native anatomy using the ‘Segment’ function of SPM12. The conventional control site of the vertex was chosen given the fact that it had been widely used in many TMS studies as the control site for comparison with the AG (e.g., Yazar *et al*., 2017; Bonnici *et al*., 2018; Branzi *et al*., 2021; Kwon *et al*., 2022). The vertex was defined using anatomy landmarks, as the midpoint between nasion and inion, along the sagittal midline (Fig. 1A).

**Figure 1.**
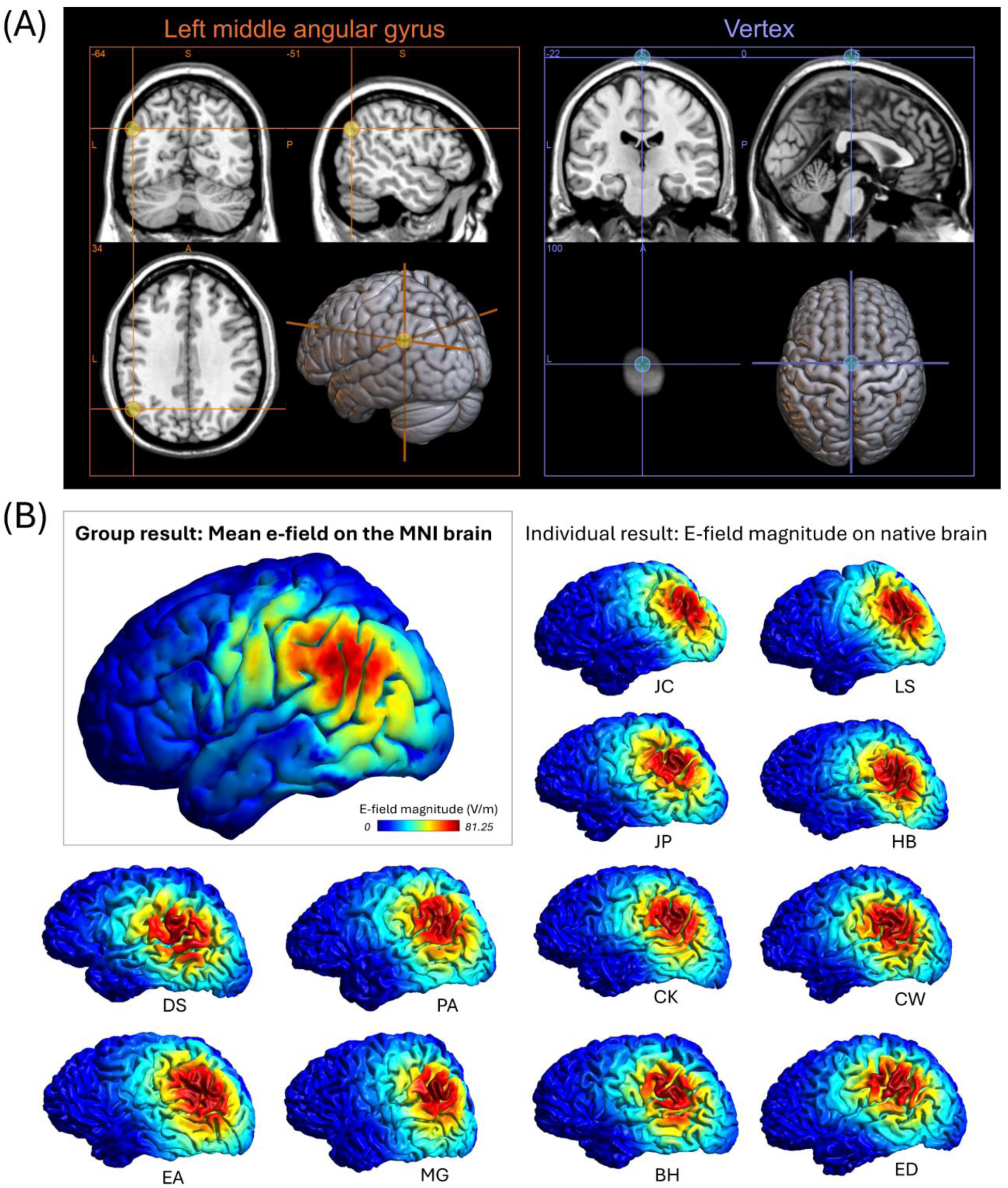
(**A**) The two targets of brain stimulation in Study 1– the left middle AG and the control site vertex – pinpointed in the standard MNI space. Note that the standard AG coordinate in the MNI space was reverse-normalised into an individual-specific corresponding coordinate in each participant’s native anatomical space as the target of brain stimulation. The vertex was also defined individually based on anatomical landmark. (**B**) The group-average result of e-field simulation (unit: volts per metre) is presented in the standard MNI template. Also presented are individual-specific e-field simulations rendered on the native anatomy of twelve representative participants.

It has been increasingly recognised that computational simulation of the TMS-induced electrical field (e-field) is important both for prospectively optimising TMS dosing and targeting (Numssen *et al*., 2024; Kuhnke *et al*., 2025) and retrospectively characterising the extent of neurostimulation (Kuhnke *et al*., 2020b). We therefore used the SimNIBS toolbox (Saturnino *et al*., 2019) to estimate the spatial distribution and strength of the induced e-field at our AG target. As illustrated in Fig. 1B, the group-average e-field map showed that the highest stimulation intensity was centred at the middle section of the AG, while the field also extended into neighbouring cortex, including the adjacent supramarginal gyrus (SMG). Thus, although the estimated TMS hotspot was focused on the AG, the observed behavioural modulation may partly reflect collateral influence on of the SMG, a region that has also been implicated in multimodal and cross-modal semantic processing (e.g., Binder *et al*., 2009; Numssen *et al*., 2021; Kuhnke *et al*., 2023a). Fig. 1B also presents the individual e-field distributions in the native anatomy of 12 representative participants.

We applied continuous theta-burst stimulation (cTBS) onto the AG and the vertex as this protocol has been demonstrated to successfully modulate performance on memory-related tasks when it was applied onto the AG (e.g., Yazar *et al*., 2014; Yazar *et al*., 2017; Bonnici *et al*., 2018; Kwon *et al*., 2022). Prior to participants starting the tasks, cTBS was delivered onto the targeted site in repeated trains of 300 bursts (3 magnetic pulses per burst; 50 Hz) with inter-train-interval of 200 ms (5 Hz); the stimulation lasted for 60 seconds, with a total number of 900 magnetic pulses. We adopted the 900-pulse cTBS protocol because it previously produced robust inhibitory effects on semantic tasks in our earlier studies targeting higher-order cortical regions (Chiou and Lambon Ralph, 2016b; Chiou and Ralph, 2018). Given that the conventional 600-pulse protocol was originally developed for the motor cortex (Huang *et al*., 2005), we reasoned that a longer stimulation duration might be more effective for modulating higher-order functions of the AG, hence our decision to adopt the 900-pulse protocol. Immediately after the cTBS, participants performed the semantic memory task, which was followed by the episodic memory task (details below). The stimulation was set at 80% of resting motor threshold (RMT), the minimum stimulation intensity on the motor cortex that caused a visible finger twitch; for testing individual RMT, we applied single-pulse stimulations to the left primary motor cortex; the value was defined as 80% of the minimum strength sufficing to trigger visible twitches in the left abductor pollicis muscle on six out of ten contiguous trials. The averaged cTBS intensity was 44.7±4.2% of the machine maximum output (range: 35% – 50%). This dosing option was based on empirical observations that RMT is measured under relaxation of fingers and generally requires stronger intensity than active motor threshold (which requires fingers pinching), hence stronger intensity for subsequent cTBS. We reasoned that RMT would provide a sufficiently strong level of TMS for modulating higher-order cortical regions such as the AG. Active cTBS was applied in both the AG session and vertex session; we did not include sham stimulation in our study. The AG and vertex sessions were, on average, 5.8±6 days apart.

Participants performed two tasks in each TMS session: a semantic memory task, followed by an episodic memory task. In the semantic memory task, participants were asked to decide whether the two words presented in each trial were semantically related to each other or not. As illustrated in Fig. 2A and 2B, concrete and abstract words were presented in separate trials (i.e., there was no mixed-category trials containing one concrete and one abstract word), but all of the trials, irrespective of being concrete or abstract, were randomly interleaved and presented in a single block of 300 trials. We designed three different conditions to probe the influences of short-term memory buffering and multimodal integration: (*i*) In the SimVis condition (the left panel in 2A and 2B), two words were presented *simultaneously* and *visually* on the screen until a response was made or up to four seconds. (*ii*) In the VisVis condition (the middle panel in 2A and 2B), a visual word (reference) was initially presented for 0.5 second, followed by a 3-second blank-screen interval during which participants had to hold the reference word in mind; subsequently, the second visual word (target) was shown and participants had up to 4 seconds to answer whether the target was semantically related to the reference word. (*iii*) In the SpchVis condition, (the right panel in 2A and 2B), participants initially heard an uttered word (reference) and had to hold the reference word in mind during the blank; when the visual target word was shown, they made a response to answer whether the reference and target were semantically related. Participants used the index and middle fingers of their right hand to indicate relatedness by pressing one of the two designated buttons. The mapping between buttons and relatedness was counterbalanced across individuals. They were told to respond as quickly and accurately as possible when the target word was shown during response window. With this design, maintaining information in a temporary buffer of short-term memory was necessary for scoring a correct answer in the VisVis and SpchVis condition whereas it was not necessary in the SimVis condition, where reference and target were presented at the same time on the screen. In addition, task-relevant information was presented solely in the visual modality in the SimVis and VisVis condition, whereas it was presented both in the auditory and visual modalities during the SpchVis condition.

**Figure 2.**
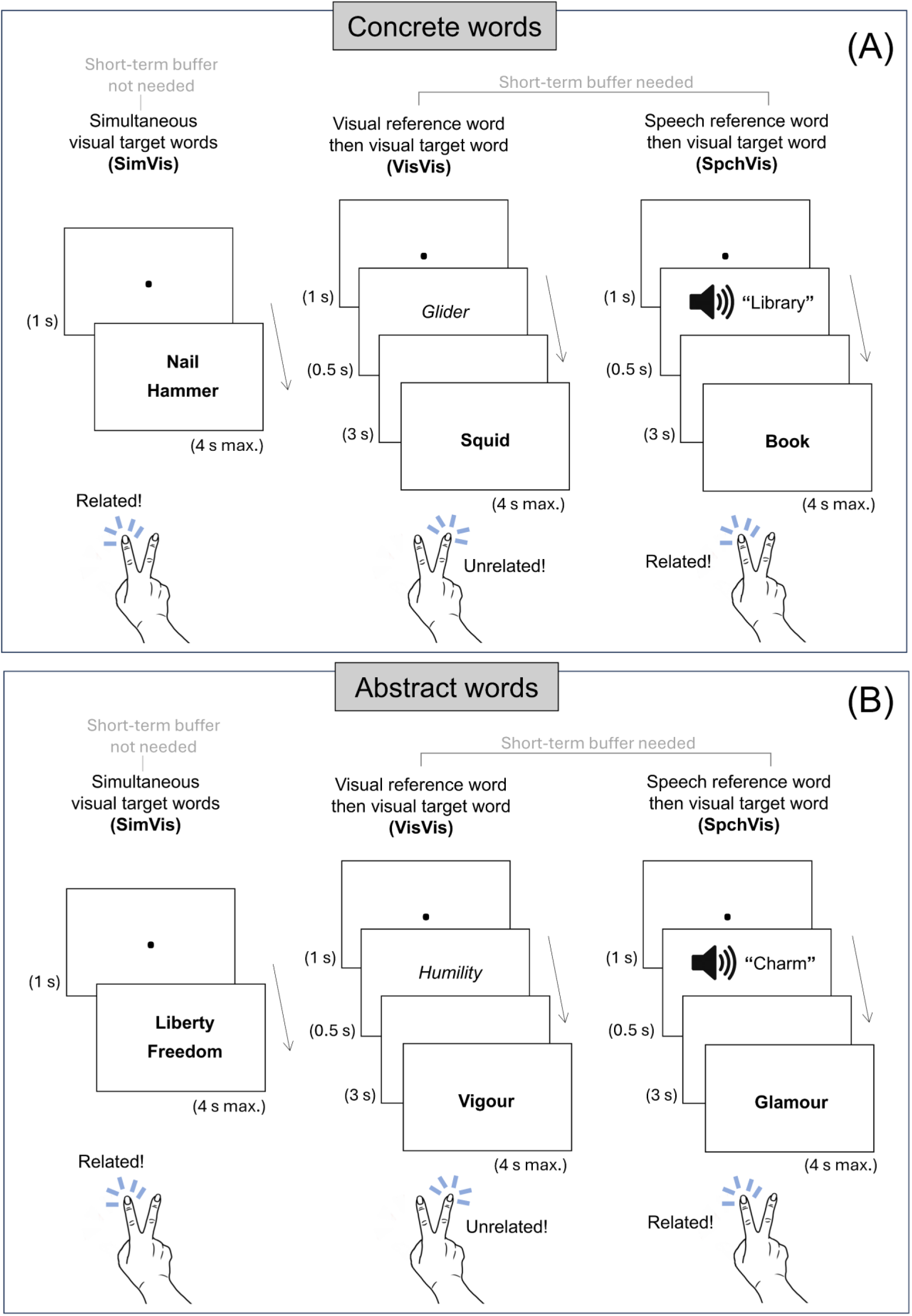
(**A**) Different task procedures when concrete words were used as stimuli. (**B**) Different task procedures when abstract words were used as stimuli. Note that all types of different trials were mixed in a single block of trials, and the mapping between buttons and relatedness was counterbalanced across participants.

We prepared a total of 960 words, comprising 480 concrete and 480 abstract items. Within each category, 160 words were designated as reference words and appeared twice during the experiment: once paired with a semantically related target (160) and once paired with an unrelated target (160). This ensured an equal number of related and unrelated trials for both concrete and abstract words while allowing reference items to be reused across conditions (but target words were never reused). Word length (indexed by letter count) was matched for concrete and abstract words, as well as between related and unrelated words (all *p*s >.22), while semantic relatedness between a reference and a target word (quantified using the *word2vec* function in MATLAB) was significantly higher in the related than unrelated condition (both *p*s < 10^-12^). In addition, we ensured that, between concrete and abstract words, there was no reliable difference in terms of word length (*p* > 0.13), the extent of semantic relatedness between words (*p* > 0.96), and lexical frequency (p > 0.66; based on the corpus statistics of Van Heuven *et al*., 2014), while concrete words had significantly higher concreteness ratings – using the corpus of Brysbaert et al. (2014) – than abstract words (*p* < 10^-12^). For each participant, 300 pairs of words were randomly selected from the full list (150 concrete and 150 abstract) and randomly allocated to different modes of presentation (50 in SimVis, 50 in VisVis, 50 in SpchVis), while we ensured no stimulus was repeated between the two TMS sessions within each participant. This “random selection and no repetition” procedure ensured that any differences in performance between presentation modes (and between TMS sessions) could not be attributed to stimulus-specific properties or residual memory from an earlier session, as the words appeared ‘novel’ in each session and were randomly chosen, apart from the necessary constraints of abstractness (concrete *vs.* abstract) and relatedness (related *vs.* unrelated). Prior to the TMS study, we tested 26 participants in a pilot behaviour study (none of them took part in the TMS experiment) using the same stimuli/paradigm. The pilot data showed the key effects of semantic processing, including effects of Concreteness, Relatedness, and Presentation Mode, thereby validating the adequacy of experimental design used in the later TMS investigation.

Immediately after participants completed the semantic memory task, they were asked to perform an unanticipated episodic memory task. In this task, participants were presented with a single block of 120 trials. On each trial, a single word was presented visually at the centre of the screen, and participants indicated whether the word was ‘old’ (i.e., encountered earlier in the semantic task) or ‘new’ (i.e., not presented previously) by pressing one of the two designated buttons. Accuracy was emphasised over speed in this task, and unlimited time to respond was allowed on each trial. The set of old items comprised 60 reference words randomly selected from each one’s individual-specific stimuli list used in the semantic task of the corresponding TMS session (either the AG or vertex), including 30 concrete and 30 abstract words. These old words were randomly intermixed with 60 new words (30 concrete and 30 abstract), yielding a total of 120 trials. The semantic task lasted for ∼50 minutes, while the episodic memory task lasted ∼15 minutes. Participants were not instructed to memorise the stimuli during the semantic task and were unaware that a subsequent memory test would follow (although the episodic memory test might be only fully unexpected the first session).

### Study 2

Full methodological details for Study 2 are reported in Chiou et al. (2025). Here we focus on the aspects of the methods relevant to the present study. In two of the conditions, we investigated the neural correlates associated with concrete and abstract words. Each trial showed a reference word above three options; all words indicated concrete concepts in the concrete blocks (e.g., reference: box; options: carton, vest, grater) while all words indicated abstract concepts in the abstract blocks (e.g., reference: onus; options: duty, affair, fame). Participants were asked to answer which of the three options was conceptually more similar to the reference. Using the corpus rating of Brysbaert et al. (2014), we ensured that words in the concrete condition were judged as more concrete than those in the abstract condition (p < 10^−25^), while word length and lexical frequency were matched between them (both *p*s > 0.45). In a pilot behavioural experiment, eight volunteers confirmed that the target and reference words were judged as conceptually more similar with each other compared with foils (*p*s < 0.001 for both concrete and abstract words). In addition to the data of concrete and abstract semantics, we also included the data of a non-semantic numerical task in which participants viewed four numbers in each trial (a 3-digit number as reference on top with three numerals below) and were asked to answer which option was numerically closer to the reference number. Each block was 19.5 seconds in duration, consisting of 5 trials; each trial contained 0.4-second fixation, followed by 3.5-second stimuli. Data from the following conditions were included in our analysis: Concrete Semantics, Abstract Semantics, Numerical Magnitude, across 10 runs of scans. In order to ensure spatial consistency across our multimethod investigation, we used the same coordinate in the MNI space that we used in the TMS experiment of Study 1 (*X* =-48, *Y* =-64, *Z* = 34) to define a spherical ROI (radius = 10 mm) centred at this coordinate, allowing direct comparison between TMS-induced behavioural effects and fMRI activity in a corresponding cortical site within the AG. As the AG is cytoarchitecturally heterogeneous (Seghier, 2013), it is important to ensure our result remains robust despite potential heterogeneity. Thus, in addition to the results of 10-mm radius ROI reported in the initial analysis, we ran verification analysis using three ROI sizes (radius: 5 mm, 15 mm, 20 mm) to ensure that conclusion would be unaffected by reasonable variation in ROI extent. Contrast parameters (Concrete > Numerical; Abstract > Numerical) were extracted from the spherical AG region and submitted to a linear mixed modelling analysis with the *lme4* package implemented in *R* (Bates *et al*., 2015).

### Study 3

Full methodological details of Study 3 are reported in Hoffman et al. (2015). Here we only describe the aspects of the methods relevant to the present study. The researchers presented concrete and abstract words in separate blocks of trials and ensured that concrete words had significantly higher concreteness and imaginability ratings than abstract words. On each trial, participants were presented with a reference word and three choices and were asked to indicate which option was most synonymous with the reference. Prior to the display of the reference with choices, two sentences were shown as a contextual cue; the meaning of the sentence cue could be related to the subsequent reference word (e.g., two sentences describing horse riding followed by a reference ‘saddle’) or unrelated to the subsequent reference (e.g., sentences about a carnival followed by a reference ‘keyboard’). The authors also included a numerical task as comparison baseline in which participants viewed four numbers in each trial (a 2-digit reference above three 2-digit choices) and were asked to answer which was numerically closer to the reference. We performed PPI connectivity analysis in SPM12: Three variables were included in the PPI model, following the standard procedure (O’Reilly *et al*., 2012) – (*i*) the estimated physiological series extracted from of the spherical AG seed (radius: 10 mm, using the same coordinate as Study 1 and 2); (*ii*) the task contrast between concrete and abstract words, which ensured that the influence of ‘concrete *vs.* abstract’ contrast was statistically excluded from the computation of PPI connectivity; (*iii*) the interaction between the AG’s timeseries and task contrast, which revealed whether there was a change in connectivity with the AG between concrete and abstract semantics. We also extracted comparison parameters (Concrete > Numerical; Abstract > Numerical) from the same spherical ROI at the AG used as the seed for PPI connectivity. It is worth noting that we chose to perform PPI analysis on the Hoffman et al. (2015) dataset because it had a task structure especially conducive for PPI. As suggested by O’Reilly et al. (2012), factorial design – e.g., the 2 (concrete *vs.* abstract) × 2 (related *vs.* unrelated) design employed by Hoffman et al. (2015) – are particularly well suited to investigating PPI connectivity. Collapsing across relatedness contexts can increase statistical power to detect contextual changes in connectivity that differed between concrete and abstract semantics. By contrast, Study 2 (Chiou *et al*., 2025) did not have a data structure amenable to PPI analysis. Following the recommendation by Coutanche and Thompson-Schill (2012), Chiou et al. (2025) employed a design with few blocks per run, distributed across ten short acquisition runs, in order to maximise between-run multivoxel decoding accuracy. Thus, this dataset (fewer blocks per run with many runs) did not provide an optimal context for PPI, which usually requires many lengthy blocks within a long run and, ideally, a factorial experimental structure. We began with a whole-brain PPI, followed by the extraction of average parameters from *a priori* ROIs to test how connectivity with the AG manifested in target regions of the semantic network. For the definition of ROIs, we used clusters from a meta-analysis on the neural substrates of semantic cognition (Jackson, 2021) to incorporate the full scope of areas. Given the regional heterogeneity within the semantic network, we subdivided the masks with the Harvard-Oxford Atlas (Desikan *et al*., 2006). The ROI was classified into two groups: (*i*) *semantic representation regions*, comprising four subparts of the anterior temporal lobe (ATL) – the superior ATL, middle ATL, inferior ATL, and temporal pole; (*ii*) *semantic control regions*, comprising three subdivisions of the left inferior frontal gyrus (IFG; anterior: *pars orbitalis*; middle: *pars triangularis*; posterior: *pars opercularis*) and one cluster in the left posterior mid-temporal gyrus (pMTG). Two versions of PPI analysis were conducted – the original analysis (Standard PPI) was conducted using the SPM12 toolbox; the replication analysis (Generalised PPI) was conducted with the gPPI toolbox (McLaren *et al*., 2012).

### Study 4

Study 4 entailed a single 1-hour behavioural testing session. We used a modified procedure of MDES that has been widely adopted to investigate thought patterns during various task situations (e.g.,Sormaz *et al*., 2018; Konu *et al*., 2021; Smallwood *et al*., 2021b; Chitiz *et al*., 2025). Participants were asked to perform a conceptual similarity judgment using the stimuli of the concrete and abstract conditions from the study of Chiou et al. (2025). The experiment comprised 200 quadruplets of words (100 concrete and 100 abstract). On each trial, a reference word was presented at the top of the screen together with three option words, and participants indicated via button press which option was conceptually most similar to the reference within the 4-second response window. A key feature of the MDES approach is that it aims to sample thought content while participants are in a relatively stable and uninterrupted cognitive state. For example, in the study of Sormaz et al. (2018), 0-back task (volunteers focused on visual stimuli) and 1-back task (volunteers focused on short-term memory) were presented in separate blocks to promote sustained engagement with external stimuli *vs.* internal memories, respectively. Guided by this rationale, in Study 4 we presented concrete and abstract stimuli in separate halves of the experiment, with the order of stimulus type counterbalanced across participants. While participants made conceptual similarity decisions, intermittent thought probes were administered: following a minimum of 20 consecutive trials (up to 25 trials), a set of 15 MDES questions was presented to query the content of their ongoing thoughts during task performance. The MDES comprised 15 questions assessing the following psychological aspects: (1) Difficulty:‘*I was thinking about how difficult the task felt*’; (2) Focus:‘*I stayed focused on answering the questions*’; (3) Future:‘*I was thinking about things that might happen in the future*’; (4) Past:‘*I was thinking about things that had happened in the past*’; (5) Self:‘*I was thinking about myself*’; (6) Others: ‘*I was thinking about other people*’; (7) Emotion: ‘*The emotional tone of my thoughts was positive (or negative)*’; (8) Images: ‘*I was visualising scenes, faces, or objects in my mind*’; (9) Details: ‘*My thoughts were detailed, specific, and defined*’; (10) Control: ‘*I maintained volitional control over my thought process*’; (11) Solution: ‘*I focused on finding the answers to the semantic task*’; (12) Intrusion: ‘*I experienced intrusive thoughts that wasn’t controllable*’; (13) Absorption: ‘*I was deeply absorbed in what I was thinking*’; (14) Distraction: ‘*My thoughts distracted me from completing the semantic task*’; (15) Words: ‘*I was thinking in words or sentences (inner speech)*’. Participants used a 10-point Likert scale to answer the questions; for all of the questions (except Question 7) the scale was “1: Not at all – 10: Completely”, and participants selected a number that best described how closely each statement matched their thoughts; for Question 7 the scale was “1: Very positive – 10: Very negative”, and participants chose a digit to indicate the valence of their emotion. Across the experiment, the MDES-question probes were administered eight times, evenly distributed in the session, with four times in the concrete condition and four in the abstract condition. Following the established method of Smallwood et al. (2021b), IBM SPSS Statistics was used to perform a principal component analysis (PCA) to extract the key dimensions that best characterised the full dataset, comprising responses to 15 questions across eight repetitions (four in the concrete condition and four in the abstract condition) concatenated across 36 participants. The default threshold (eigenvalue > 1) of SPSS was used to identify components; split-half reliability index with Spearman-Brown correction (Eisinga *et al*., 2013) was calculated to ensure the robustness of each identified component. The scree plot and the percentage of variance of each component are reported in Supplemental Fig. 1.

### Study 5

Full methodological details are reported Chiou et al. (2020). Here we focus only on the aspects relevant to the re-analysis. We extracted AG activity from the same AG region used throughout the present investigation – sphere centred at *X* =-48, *Y* =-64, *Z* = 34 (radius: 10 mm), using the data of a social-semantic task of Chiou et al. (2020). In each trial of this task, participants read an adjective describing a personality trait (e.g., conscientious, stubborn, optimistic, etc.). In the blocks of the Self condition, they answered whether the adjective suitably described their own personality, whereas in the blocks of the Queen condition, they answered whether the adjective was consistent with their understanding about the personality of Queen Elizabeth II. A total of 180 adjectives were used, with 90 assigned to either the Self or Queen condition and counterbalanced across participants to minimise stimulus-driven effects. Each block consisted of five trials; each trial began with a fixation dot (0.8 second) followed by a word (2.8 seconds). In addition to the social-semantic task, there was a non-semantic perceptual task: In this task, participants viewed meaningless patterns (squiggly lines created from scrambling the text used in the Self and Queen condition) and decided whether the two patterns of each trial were left/right flipped. Contrast parameters were extracted from the AG to examine whether its activation differed when semantic processing was compared against a minimally semantic baseline (the perceptual task) versus a richly semantic baseline (passive rest). We also performed the same analysis on the data of Chiou et al. (2025) and Hoffman et al. (2015) to examine the influence of semantically impoverished baseline (numerical task) versus semantically rich baseline (rest) on the polarity and magnitude of AG activation.

## Results

We begin by previewing our key findings to highlight the functional profile of the AG. In Study 1, cTBS to the AG impaired semantic performance when information had to be temporarily buffered, with a significantly greater disruption for abstract than concrete semantics. This pattern indicates that the AG is especially engaged when abstract-semantic information needs to be transiently cached in an online buffer. In Study 2, we found that when the influence of cognitive effort (quantified by reaction times) was statistically partial out, concrete semantics no longer induced higher AG activity than abstract semantics, suggesting that the apparent concreteness-related AG effects might be substantially confounded with non-semantic mental activity during a cognitively less effortful situation. In Study 3, we found that while the AG was significantly less active for abstract than concrete semantics (replicating Study 2), there was enhanced connectivity between the AG and several regions involved in the controlled retrieval of semantic knowledge. In Study 4, MDES was used to examine how thought patterns differ during the processing of concrete and abstract semantics. We found that, relative to abstract semantics, participants reported being more absorbed in visual imagery and experiencing less mental effort (implying automaticity) during the processing of concrete semantics. Finally, in Study 5, we showed that AG activation was highly contingent on the choice of comparison baseline. When semantic tasks were contrasted with semantically impoverished perceptual or numerical tasks, the AG appeared *positively engaged*. However, when contrasted against semantically rich passive rest, the same data yielded markedly reduced responses or even *negative deactivation*. These findings underscore that interpretation of AG involvement must be made with caution, as baseline selection can fundamentally alter the apparent direction and magnitude of activation. We also discussed the utility of rest as a baseline.

### Study 1

For the reaction time data (RT) of the semantic task, prior to any statistical analysis, we excluded trials with RTs faster than 100 ms or exceeding three standard deviations above the condition mean and included only trials with correct responses. Across participants, outlier RTs accounted for less than 0.5% of trials. The RT and accuracy of every condition are reported in Supplemental Table 1. None of the participants were excluded. RT data were analysed using a repeated-measure ANOVA, with within-participant factors of TMS (AG *vs.* vertex), Concreteness (concrete *vs.* abstract), Relatedness (related *vs.* unrelated), and Presentation Modes (SimVis, VisVis, and SpchVis). We first report statistics pertinent to our aim of assessing whether disruption of the AG would have any causal consequence on behavioural performance in the semantic task. There was a significant main effect of TMS (*F*_(1, 23)_ = 13.91, *p* = 0.001, Fig. 3A), with stimulation to the left AG yielding slower responses (1213 ms) than cTBS of the control site (vertex; 1108 ms). Crucially, the statistics also identified a significant interaction between TMS and Concreteness (*F*_(1, 23)_ = 6.19, *p* = 0.02, Fig. 3B). Post-hoc comparisons were performed to identify the source of this interaction effect: Although the influence of cTBS to the AG (quantified as AG *minus* vertex) was significant for both concrete (*p* = 0.004) and abstract semantics (*p* < 0.001), the overall magnitude of cTBS influence was reliably larger under the condition of abstract words (130 ms) as compared to concrete words (78 ms, *t*_(23)_ = 2.48, *p* = 0.01), suggesting that although the semantic processing on both types of words was negatively affected by cTBS to the AG, the disruptive effect was disproportionately greater for abstract than for concrete semantics. Moreover, we also found a strong trend indicating an interaction between TMS and Presentation Modes (*F*_(1, 23)_ = 3.01, *p* = 0.06, Fig. 3C). Therefore, Post-hoc comparisons were conducted on an exploratory basis to examine the pattern behind this interaction trend: While the cTBS impact on the AG (as compared to the vertex) was significant in all three kinds of Presentation Modes (SimVis: *p* = 0.02; VisVis: *p* < 0.001; SpchVis: *p* < 0.001), the effect was reliably smaller during the SimVis condition which did not require information to be maintained in a memory buffer, compared with the VisVis and SpchVis conditions that both required online buffering (SimVis *vs.* VisVis: *t*_(23)_ =-1.77, *p* = 0.04; SimVis *vs.* SpchVis: *t*_(23)_ = - 2.28, *p* = 0.01; no difference between VisVis and SpchVis: *t*_(23)_ =-0.24, *p* = 0.41). These findings suggest that buffering demands may be a modulator of AG engagement in semantic tasks, with greater reliance on AG neural resources when active maintenance is required.

**Figure 3.**
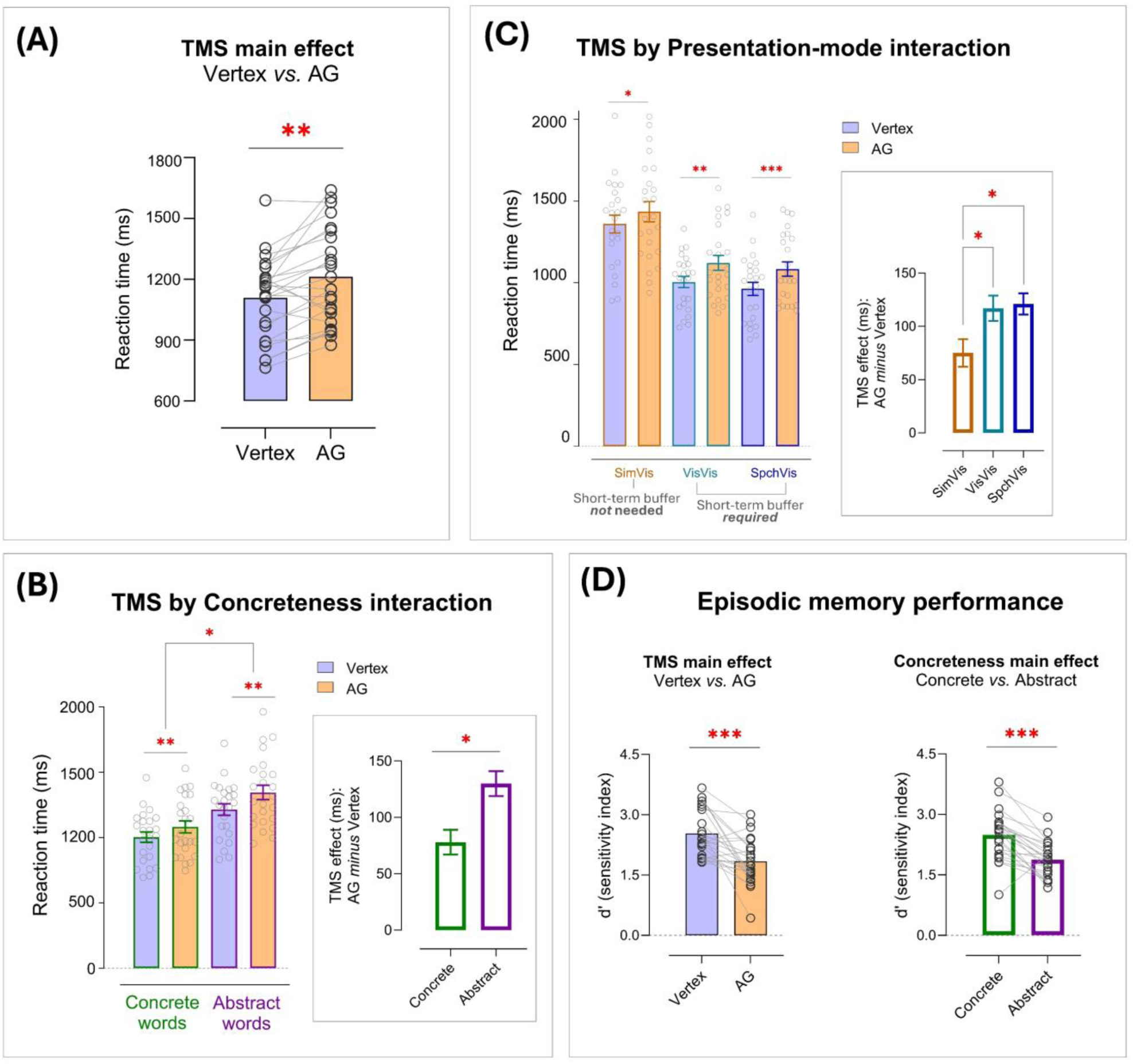
(**A**) The main effect of TMS (AG *vs.* Vertex) on RT. The circles indicate individual datapoints. ** *p* = 0.001; (**B**) The interaction between TMS (AG *vs.* Vertex) and Concreteness (concrete *vs.* abstract). The circles indicate individual datapoints. Error bars indicate standard error of the mean (SEM). * *p* < 0.05; ** *p* < 0.01; (**C**) The interaction between TMS (AG *vs.* Vertex) and Presentation-mode (SimVis *vs.* VisVis *vs.* SpchVis). The circles indicate individual datapoints. Error bars indicate standard error of the mean (SEM). * *p* < 0.05; ** *p* < 0.01; *** *p* < 0.001; (**D**) Results of the episodic memory experiment. (left) The main effect of TMS on d-prime. (right) The main effect of Concreteness on d-prime. The circles indicate individual datapoints. *** *p* < 0.001

For completeness, we also report other significant statistics in the RT analysis not involving the factor of TMS. We found main effects of Concreteness (*F*_(1, 23)_ = 199.12, *p* < 0.001), Relatedness (*F*_(1, 23)_ = 33.88, *p* < 0.001), and Presentation Modes (*F*_(1, 23)_ = 155.63, *p* < 0.001). We also found interactions between Concreteness and Presentation Modes (*F*_(1, 23)_ = 6.21, *p* = 0.004) and between Relatedness and Presentation Modes (*F*_(1, 23)_ = 30.07, *p <* 0.001). For accuracy, there were no significant effects involving the TMS factor (all *p*s > 0.13), while there were significant main effects of Concreteness (*F*_(1, 23)_ = 104.66, *p* < 0.001), Relatedness (*F*_(1, 23)_ = 44.45.12, *p* < 0.001), as well as interactions – Concreteness × Relatedness (*F*_(1, 23)_ = 22.49, *p* < 0.001), Relatedness × Presentation Modes (*F*_(1, 23)_ = 11.50, *p* < 0.001), and Concreteness × Relatedness × Presentation Modes (*F*_(1, 23)_ = 4.33, *p* = 0.02).

Given the substantial inter-individual variability common in TMS research, we additionally analysed RTs using linear mixed-effects modelling, allowing the effects of the task variables to vary across participants through random slopes. Session (first *vs.* second) was also included to explain potential order effects. As detailed in Supplemental Tables 2 and 3, we constructed a series of models with different random-effect structures and compared their fit to data using AIC and BIC. This procedure identified the model with *random slopes for all variables* as the best-fitting model. Subsequent test for this winner model yielded a pattern broadly consistent with the ANOVA results, including all the principal effects involving TMS: (*i*) a significant main effect of TMS, *F*_(1, 10.79)_ = 17.00, *p* = 0.002; (*ii*) a significant TMS × Concreteness interaction, *F*_(1, 446.13)_ = 8.83, *p* = 0.003; post-hoc comparisons showed that TMS effects were larger for abstract words, *t*_(29.1)_ = 4.20, *p* = 0.0002, than concrete words, *t*_(29.1)_ = 2.45, *p* = 0.02; (*iii*) a trend towards a TMS × Presentation interaction, *F*_(2, 446.13)_ = 2.74, *p* = 0.066; post-hoc tests showed that TMS effects were larger in the two contexts requiring information-buffering (SpchVis: *t*_(36.3)_ = 3.72, *p* = 0.0007; VisVis: *t*_(36.3)_ = 3.58, *p* = 0.001) relative to the non-buffering context (SimVis: *t*_(36.3)_ = 2.26, *p* = 0.03).

Although previous TMS studies have examined the contribution of the AG to episodic memory, most have focused on its role in recollection *after* the content has been encoded already (e.g., Yazar *et al*., 2014; Yazar *et al*., 2017). In contrast, the present study applied TMS *prior to* encoding, allowing us to test whether AG disruption impairs subsequent encoding and, in turn, later retrieval. For the episodic memory data, we analysed the results of both d-prime values (*d*’, defined as the difference between hit rate and false alarm rate: *Z*_(Hit)_ − *Z*_(F.A.)_) and accuracy rates. As the two sets of analyses yielded entirely consistent results, here we only report the *d*’ results: we submitted the data to a repeated-measure ANOVA, with two within participant factors of TMS (AG *vs.* vertex) and Concreteness (concrete *vs.* abstract). Results revealed two significant main effects. As revealed in Fig. 3D (left), following AG stimulation, participants recalled significantly less memory relative to their performance after vertex stimulation (*F*_(1, 23)_ = 21.61, *p* < 0.001). We also found that although the proportion of Hit responses significantly declined following AG stimulation as compared to vertex stimulation (*t*_(23)_ =-6.18, *p* < 0.001), the proportion of False Alarm responses was unaffected (*t < 1, p* > 0.53), suggesting that AG stimulation reduced the amount of information successfully encoded (thus later recalled), rather than causing confusion in discerning new from old. In addition, as shown in Fig. 3D (right), participants retrieved significantly fewer abstract words as compared to concrete words (*F*_(1, 23)_ = 35.94, *p* < 0.001). The interaction between TMS and Concreteness was not significant (*F* < 1, *p* > 0.69). Taken together, these results replicate the well-documented concreteness effect in the memory recollection literature (e.g., Paivio *et al*., 1994) and further demonstrate that disrupting the AG impairs memory encoding, thereby reducing subsequent recall.

### Study 2

In Study 2, we re-analysed the fMRI data from Chiou et al. (2025). To ensure comparability across all TMS and fMRI datasets, the same AG coordinate, stimulated in the TMS experiment (Study 1), was used to define a spherical ROI (10mm radius) for all our fMRI analyses. As shown in Fig. 4A, when semantic tasks were contrasted against a numerical task as the baseline, only concrete words elicited significantly above-baseline AG activity (*p* = 0.04). Activation for abstract words was not significantly above baseline (*p* = 0.22), and AG activity was robustly greater for concrete than abstract semantics (*t*_(24)_ = 4.96, *p* < 0.001). While this pattern replicated the concreteness effect in the literature (Hoffman and Bair, 2025), it stood in contrast to our TMS findings (Study 1), in which AG disruption disproportionately impaired abstract words. To clarify this discrepancy, we examined behavioural performance of the two conditions and found a highly robust difference, with responses in the concrete condition being reliably faster (*t*_(24)_ =-12.45, *p* < 0.001, Fig. 4B) and more accurate (*t*_(24)_ = 7.70, *p* < 0.001) than in the abstract condition. These results raise the possibility that the concreteness effect in AG activity might reflect differential cognitive effort between conditions, rather than a true preference for concrete over abstract semantics, with increased effort associated with reduced AG engagement. This interpretation is consistent with evidence linking the AG to the default mode network, which typically exhibits task-related *deactivation* under higher demand, externally-oriented focus on perceptual stimuli and time-pressured tasks (Humphreys *et al*., 2015; Murphy *et al*., 2019; Kuhnke *et al*., 2023b).

**Figure 4.**
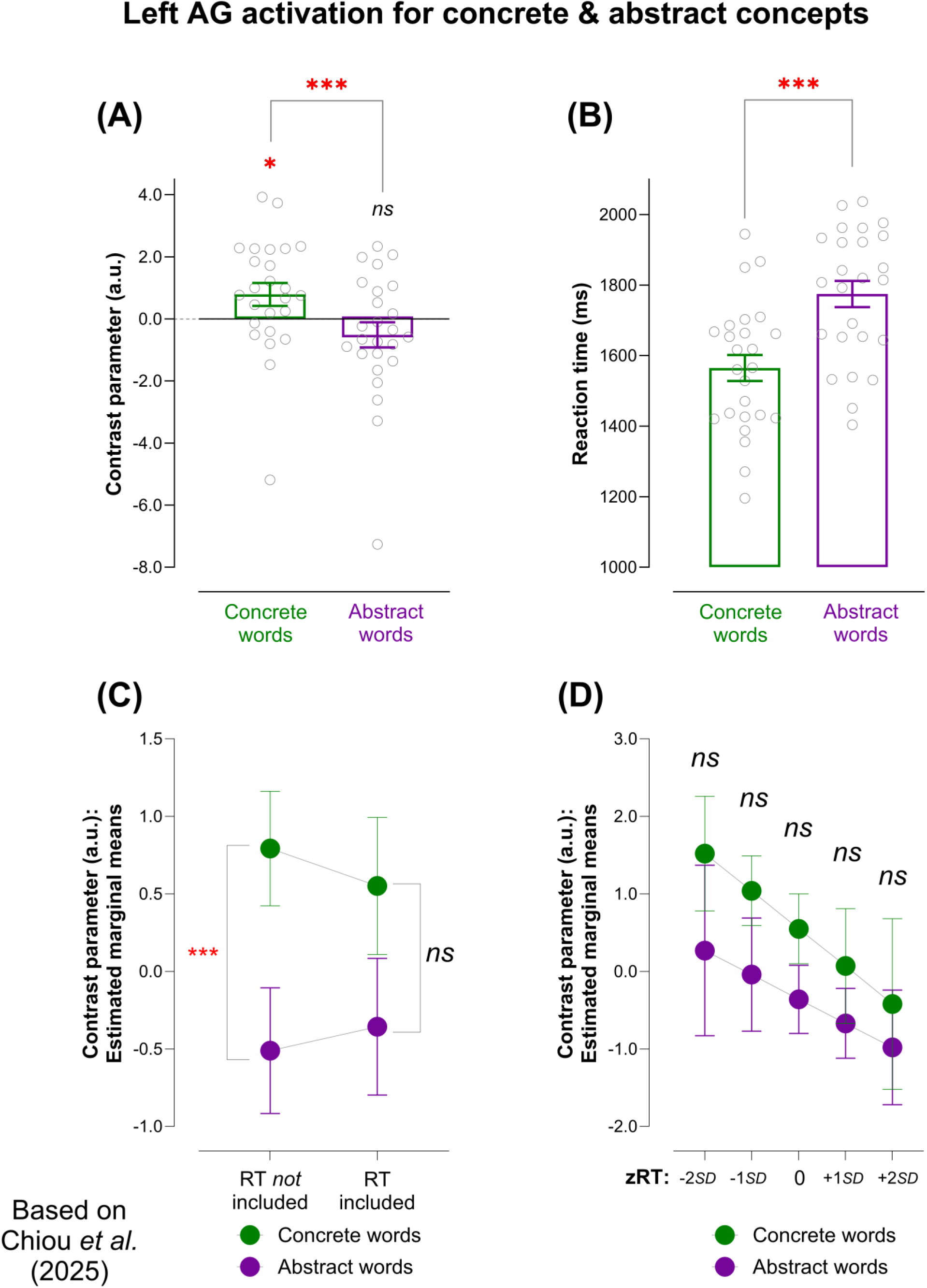
These results are based on the data of Chiou et al. (2025). (**A**) The AG was significantly more active for concrete than abstract semantics. (**B**) Behavioural RT was significantly faster for concrete than abstract words. (**C**) Using linear mixed-effect modelling, we found that the concreteness effect disappeared when RT was included as a co-variate. We also found that the concreteness effect was eliminated when another effort-sensitive index – accuracy – was included in the model but this effect persisted when participants’ age was included (see text). (**D**) AG activity declined as RT prolonged and, critically, there was no concreteness effect at any level of RT. * *p* < 0.05; *** *p* < 0.001

To clarify how cognitive effort modulates AG activity and interacts with semantic content, we used linear mixed-effects modelling incorporating RT as a proxy for cognitive effort (model formulae are in Supplemental Table 4). Because the inclusion of RT changed the concreteness effect from highly significant to non-significant, we performed Bayesian statistics to quantify how the weight/polarity of evidence shifted before and after RT was added, allowing a principled interpretation of the result. As shown in Fig. 4C, when RT was not included in the model, semantic concreteness had a potent effect on AG activity (*t*_(24)_ =-4.96, *p* < 0.001, B_10_ = 545.66, which suggests overwhelming evidence favouring H_1_), with concrete words eliciting reliably greater AG activation than did abstract words. However, after RT was added to the model, semantic concreteness no longer modulated AG activity (*t*_(23.45)_ =-0.91, *p* = 0.37, B_10_ = 0.31, indicating a moderate support for H_0_/no difference). Moreover, Fig. 4D shows that AG activity was robustly modulated by RT (*t*_(35.42)_ =-4.75, *p* < 0.001, B_10_ = 334.40 – strong support for H_1_), with slower RT (more effort) associated with weaker AG activity. Crucially, controlling for RT substantively attenuated the concreteness effect such that AG activity did not differ between the two semantic conditions at any level of performance speed (all *p* > 0.07, B_10_ ranged from 0.26 to 0.93, indicating anecdotal/moderate support for no difference between concrete and abstract). Furthermore, we calculated semi-partial R^2^ to evaluate the unique variance exclusively explained by the regressor of concreteness when other factors had been partial out: When concreteness was the only regressor in the model, its unique variance was 11.3%; however, when RT was added as a co-variate, the unique variance of semantic concreteness dropped to 4.5%; when both RT and the interaction between RT and concreteness were included, R^2^ further declined to 0.5%. Together, these results mean that RT shares variance with semantic concreteness, thus undermining the explanatory power of concreteness when RT is incorporated in the equation.

To ensure whether the effect of RT can generalise to other measures that also gauge cognitive effort, we reran linear mixed-effect modelling using accuracy values as a co-variate, given the fact that processing concrete words was robustly more accurate than abstract words. Results replicated the pattern observed with RT as a co-variate: When accuracy was included in the model, the effect of semantic concreteness disappeared (*t*_(27.51)_ =-0.21, *p* = 0.84, B_10_ = 0.22 – moderate support for H_0_). We also ran a control analysis using participants’ age as a co-variate to ensure that the effect of semantic concreteness does not always reduce to non-significance whenever any co-variate is added (even when it is not an index of cognitive effort). We found that semantic concreteness remained highly significant despite the addition of age as a co-variate (*t*_(23)_ =-3.37, *p* = 0.002, B_10_ = 15.42 – strong support for H_1_), thus ruling out non-specific explanation. Together, these analyses suggest that the apparently robust concreteness effect in the AG might reflect this region’s preferential response to certain mental activities associated with cognitively less effortful states (e.g., mind-wandering, episodic recall, automaticity, imagery – which we explored in Study 4), rather than a genuine preference for concrete over abstract semantics. When cognitive effort is controlled for, semantic concreteness no longer modulates AG activity, turning the polarity of Bayes factors from overwhelming support for the existence of concreteness effect to moderate support for a null effect.

Finally, for the data of Study 2, we examined whether the pattern of results remained reliable with different ROI extents, given the high cytoarchitectural heterogeneity within the AG (Seghier, 2013). We tested three additional ROI radii: 5mm, 15mm, 20mm. As shown in Table 1, across all analyses, we identified highly significant concreteness effects before the incorporation of RT (-3.79 < *t* < - 4.97; 32.84 < B_10_ < 493.53; *p* ≤ 0.001 for all tests). However, the addition of RT substantively weakened the effect (-0.87 < *t* <-0.61; 0.25 < B_10_ < 0.31; *p* > 0.37 for all tests), supporting the reliability of our conclusion across reasonable variation in ROI scopes.

**Table 1.**
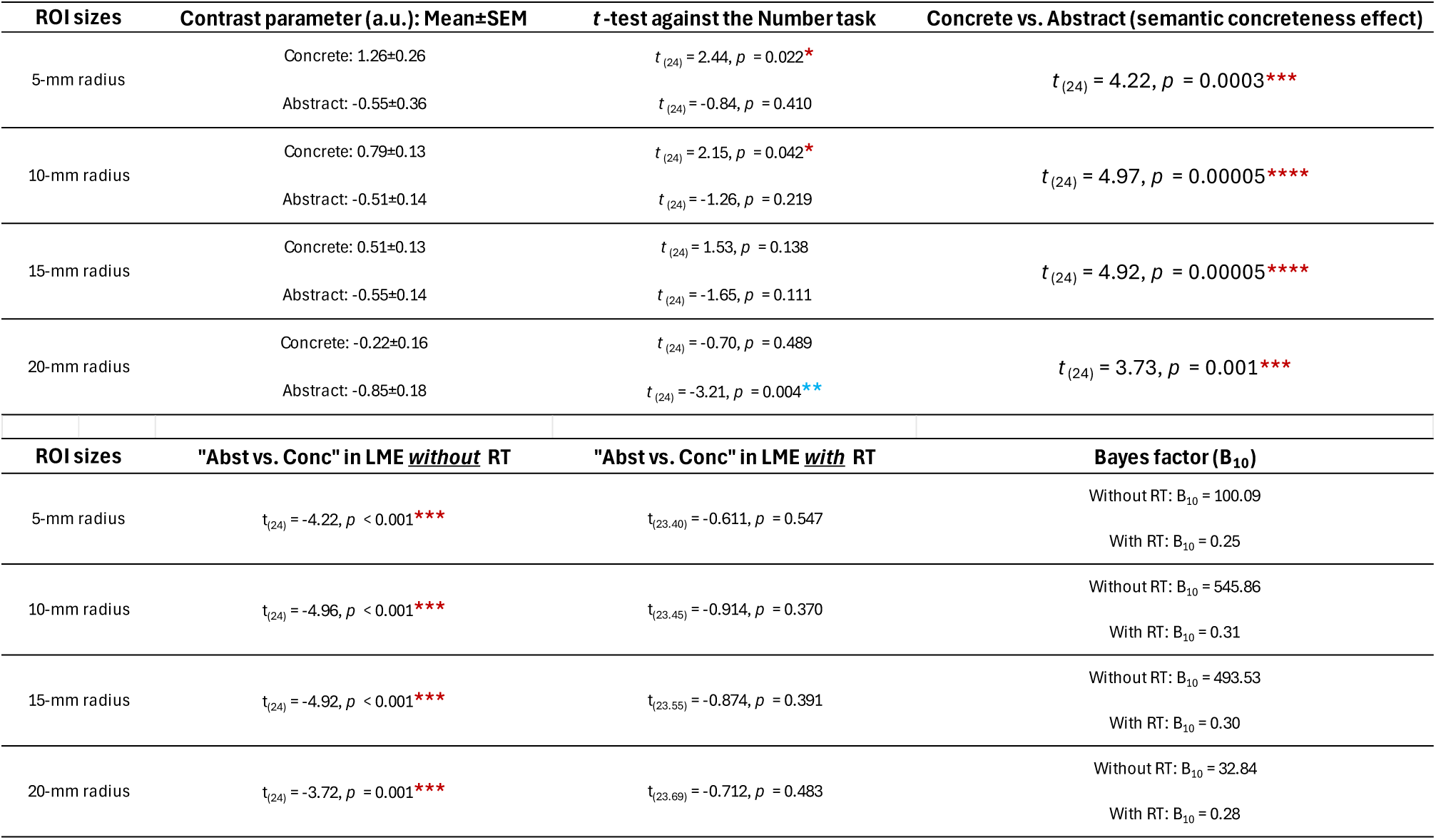
Testing different radius sizes (5mm, 10mm, 15mm, 20mm) of spherical ROI from the fMRI data of Study 2.

### Study 3

Concrete and abstract semantic tasks engage partially distinct brain networks (Hoffman & Bair, 2025): concrete semantics typically elicits stronger activation in ‘core’ nodes of the default mode network, including the left AG, medial prefrontal cortex, and posterior cingulate cortex (Shao *et al*., 2024), whereas abstract semantics reliably suppresses AG activity (relative to concrete words) while preferentially recruiting regions implicated in goal-directed control over semantic processing – the so-called semantic control network (Jackson, 2021), encompassing the left IFG and the pMTG, alongside superior portions of the ATL, a key region for abstract semantic processing (Rice *et al*., 2018). This apparent dissociation raises a key question: Although the AG shows reduced activation during abstract semantic processing, does it therefore disengage from the broader semantic network, or does it nonetheless stay coupled to regions in the semantic network despite local deactivation? Notably, reduced regional activity does not imply functional detachment, as regions can remain dynamically coupled with task-relevant networks even when exhibiting decreased activation (Krieger-Redwood *et al*., 2016; Wang *et al*., 2021). Accordingly, we re-analysed the fMRI dataset from Hoffman et al. (2015) to examine whether AG connectivity during abstract semantics preferentially links with regions for semantic representation (ATL), semantic control (IFG and pMTG), or both. Also, given variation within semantic regions, including a superior–inferior divide in the ATL (superior-verbal vs. inferior-visual; Lambon Ralph *et al*., 2017) and an anterior–posterior divide in the IFG (anterior-semantic vs. posterior-phonological; Klaus and Hartwigsen, 2019), the ATL and IFG were further segregated into subregions to assess finer-grained effects.

We first checked if in this data we could still see AG deactivation for abstract words. As shown in Fig. 5A, there was a significant concreteness effect such that AG activity for abstract semantics was reliably weaker than concrete semantics (*t*_(17)_ =-2.25, *p* = 0.03), replicating this classic effect. Having established AG deactivation for abstract semantics, we used this region as the seed of PPI to know how its pattern of connectivity altered between concrete and abstract semantics. Particularly, we tested whether PPI effects would manifest in semantic-related regions. As shown in Fig. 5B, results showed that the AG increased connectivity with two of the four ‘semantic control’ regions during the processing of abstract words (anterior IFG/*pars orbitalis*: *t*_(17)_ = 2.52, *p* = 0.01; pMTG: *t*_(17)_ = 2.44, *p* = 0.01); the effects were not significant in all ATL subregions (*p*s > 0.12). In Fig. 5B, we also reported an exploratory whole-brain interrogation to examine whether PPI effects extended beyond the semantic network. The results corroborated the pattern of ROIs findings – during the processing of abstract words, the AG was more connected with the left IFG, pMTG, and part of the superior temporal gyrus. In addition, we conducted a replication analysis using the gPPI approach (McLaren *et al*., 2012), which provides a flexible framework for modelling connectivity across all task conditions. The results were largely consistent with the standard PPI analysis implemented in SPM12, including stronger AG connectivity with the left IFG and pMTG during abstract semantic processing (see Supplemental Table 5 for peak coordinates). Together, these results caution against equating deactivation with a lack of functional involvement or withdrawal from task processing: Despite lower activation for abstract words, the AG remains integrated with the semantic network.

**Figure 5.**
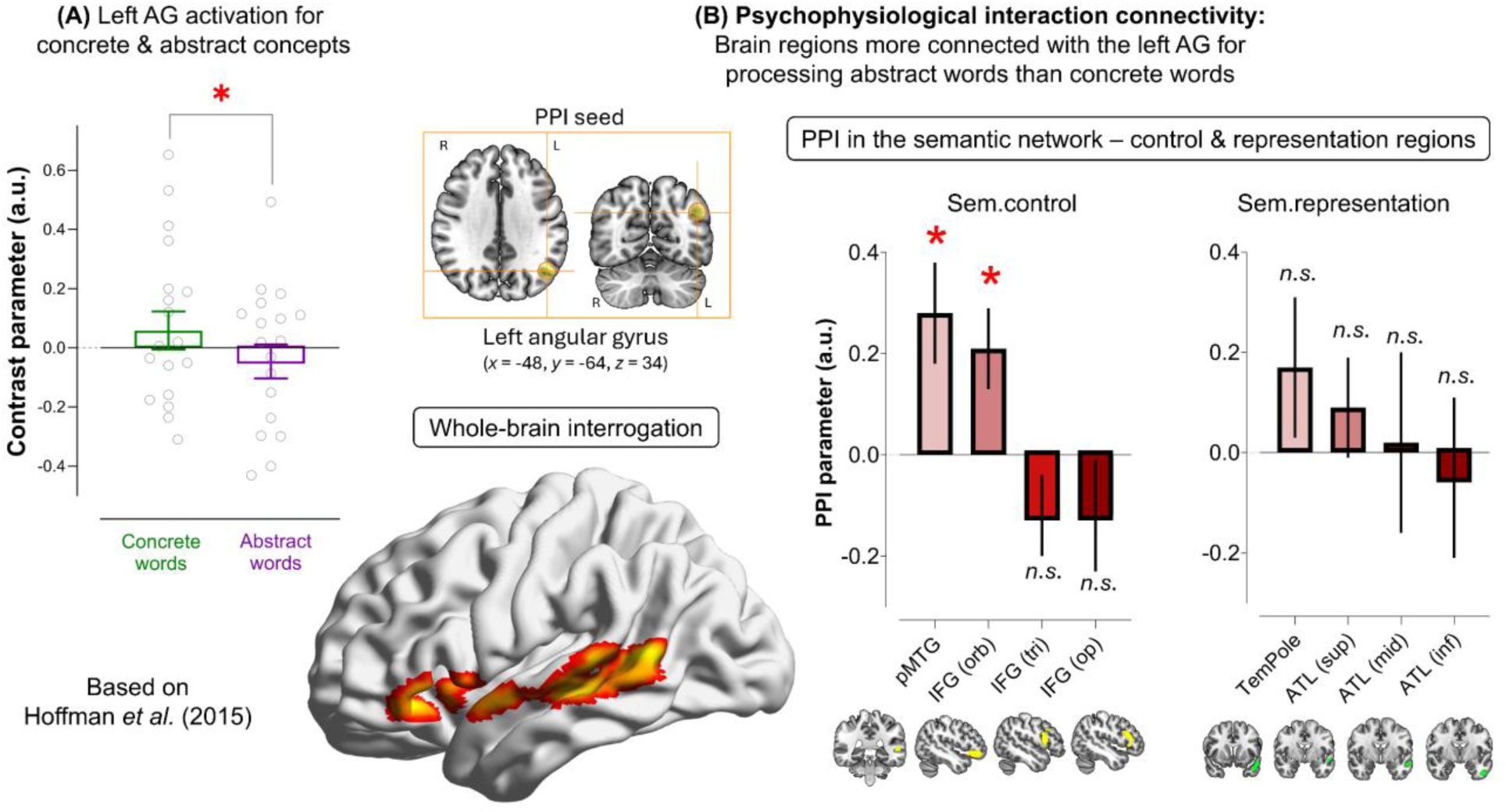
These results are based on the data of Hoffman et al. (2015). (**A**) The AG was significantly more active for concrete than abstract semantics. (**B**) The locus of AG seed for PPI connectivity was situated at *X* =-48, *Y* =-64, *Z* = 34, highlighted in the yellow circle in the inset box. Below the inset box shows significant PPI results of whole-brain search (see Supplemental Table 5 for peak coordinates). PPI effects were also examined in *a priori* ROIs in the semantic network. We found that, despite the AG being less active for abstract semantics, PPI connectivity revealed that the AG was significantly more connected with two regions crucial for semantic control (the IFG and pMTG) during the processing of abstract words. Whole-brain exploratory search was thresholded at *p* < 0.005 at the voxel level and *p* < 0.05 at the cluster level to control for multiple comparisons. Error bars indicate SEM. * *p* < 0.05 Benjamini-Hochberg FDR methods were used to correct for multiple comparisons.

### Study 4

Under a lower-demand context (e.g., the processing of concrete words), elevated AG activity is inherently ambiguous: it may reflect representations of concrete semantics, cognitive automaticity, task-unrelated thoughts, or mental imagery, as all of these have been found to engage the AG. Consequently, AG activation cannot be interpreted as only reflecting concrete semantics. To clarify the phenomenological factors that underlie increased AG activation during concrete words, we used MDES to clarify participants’ patterns of thought when they were making semantic decisions on concrete and abstract words. The procedure of our MDES experiment produced two sets of data: (*i*) the RT and accuracy rates of semantic decision and (*ii*) experience-sampling responses obtained during task performance, which were then analysed using PCA to extract components capturing participants’ thought patterns.

As predicted, we found that reaction time was faster for concrete (1330 ms) than abstract words (1731 ms; *t*_(35)_ =-16.21, *p* < 0.001) and accuracy was higher for concrete (94%) than abstract words (76%; *t*_(35)_ = 11.99, *p* < 0.001). In light of this pronounced behavioural difference, we next examined whether patterns of ongoing thoughts differed between concrete and abstract processing, by comparing PCA-derived loading scores across the two conditions. Four components emerged from the PCA: the first characterised off-task thoughts (split-half reliability: *r* = 0.95); the second captured mental absorption in visual imagery (*r* = 0.80); the third indexed the deployment of cognitive effort (*r* = 0.70); the fourth denoted the perceptual modalities of experience (*r* = 0.41). After the identification of these four statistically reliable components, we examined whether their component scores differed between concrete and abstract words. As shown in Fig. 6, we found that processing concrete words elicited significantly greater visual imagery than abstract words (*t*_(35)_ = 2.68, *p* = 0.01). In addition, participants also reported significantly less cognitive effort on the task for concrete than abstract words (*t*_(35)_ =-8.97, *p* < 0.001), suggesting automaticity or spontaneity during the processing of concrete words (as opposed to cognitive effort or controlled processes for abstract words). The two types of words did not differ on the dimensions of off-task thoughts and perceptual modalities (both *p*s > 0.28). Together, the PCA results helped elucidate the experience-related factors that underpin greater AG activity for concrete words – imagery and automaticity. These factors accord with prior evidence showing that the AG, as part of the default mode network, is closely associated with imagery/simulation (e.g., Boccia *et al*., 2015; Spreng *et al*., 2018) and automatic processing (e.g., Davey *et al*., 2015; Vatansever *et al*., 2017).

**Figure 6.**
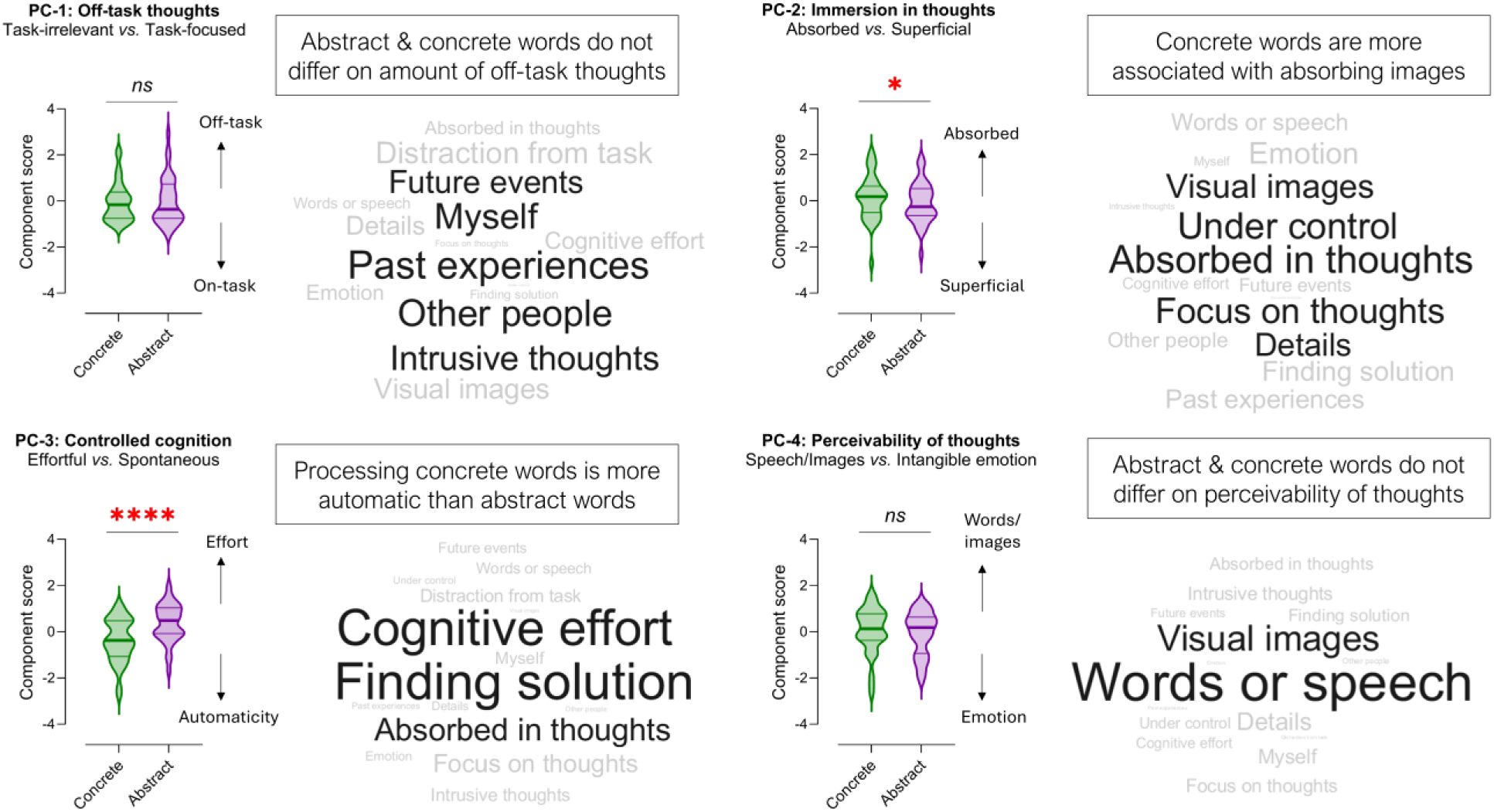
MDES revealed four statistically reliable components in the experiential sampling data. We compared the component loading scores between the concrete and abstract condition. Results showed two significant differences between the two conditions – participants reported being significantly more absorbed by visual imagery during the processing of concrete words; they also reported more cognitive effort when processing abstract words. The wordclouds visualise how each of the four components load on the 15 experiential dimensions used in MDES, with bigger font size indicating greater loading. * *p* < 0.05; **** *p* < 0.0001

### Study 5

Although passive rest is commonly adopted as a baseline in studies of AG function, it is far from cognitively blank or neutral. Moments of rest are characterised by different forms of semantically rich mentation, including autobiographical recollection, future simulation, and associative thoughts (Smallwood *et al*., 2021b; Fernandino and Binder, 2024). As a result, contrasting a semantic task against rest can partially cancel out semantic processes or even yield apparent deactivation because resting periods might contain abundant semantic content. At the same time, rest retains important methodological value: because it is difficult to design an equivalent non-semantic control across different domains and experiments, passive rest provides a practical *common reference point* for determining whether a task drives activity above or below an internally oriented mental state (Humphreys and Tibon, 2023). Interpreted carefully, the polarity of activation relative to rest can offer insight into the extent to which a given process amplifies or attenuates AG engagement beyond spontaneous mentation during rest. Accordingly, we compared semantic conditions against (*i)* non-semantic tasks and (*ii*) passive rest, to determine how baseline choice alters the patterns of AG activity and discuss the factors that underlie polarity shifts above and below the rest baseline.

We examined the effect of baseline selection across different datasets. As shown in Fig. 7A (left), when compared to a numerical task, the two semantic conditions of Chiou et al. (2025) were either weakly greater than number processing (Concrete *vs.* Number: *p* = 0.04) or no difference (Abstract *vs.* Number: *p* = 0.22). However, when the baseline was rest, both concrete and abstract conditions, as well as the number task, produced *negative deactivation* of substantial magnitude (Fig. 7A right; all *p*s < 0.0001), indicating that the ‘semantic > non-semantic’ contrast likely captures attenuation of deactivation in the semantic contexts, rather than true activation over and above the resting baseline. Put differently, semantic processing in this experiment may be insufficient to amplify AG activity beyond spontaneous semantic thoughts at rest. A similar pattern was observed in Hoffman et al. (2015): Highly significant deactivation was found when the semantic conditions – concrete and abstract – were contrasted with passive rest (both *p*s < 0.0001). Interestingly, some semantic tasks – such as the social semantic paradigm used in Chiou et al. (2020) – elicited robust AG activation regardless of whether the baseline was impoverished or abundant in semantic content. In this task, participants viewed personality descriptions and assessed whether the adjectives suitably described the character of themselves or Queen Elisabeth II. As shown in Fig. 7B (left), when compared against a non-semantic perceptual task, both the Self and Queen conditions elicited highly significant AG activation (both *p*s < 0.0001). Crucially, when compared against the resting baseline (Fig. 7 right), both conditions of this social semantic task still drove AG activation (Self: *p* = 0.004; Queen: *p* = 0.01), despite the resting baseline already containing rich semantic mentation, indicating that social cognition processing amplified AG engagement beyond the resting state. In addition to the results based on contrast parameters reported here, in Supplemental Fig. 2 we report the data based on the percent of signal change. The conclusions were identical between the two measures.

**Figure 7.**
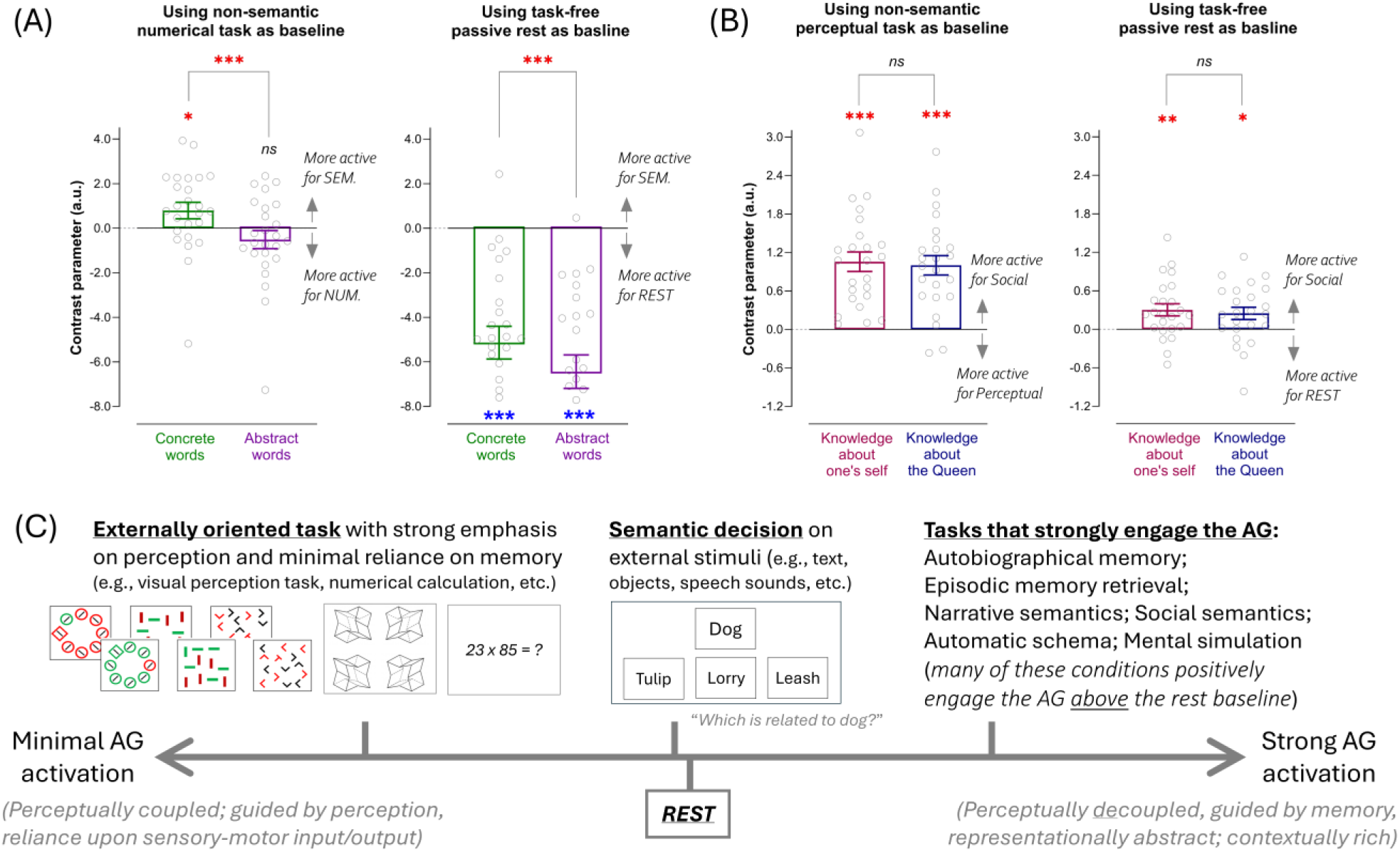
(**A**) The result was based on Chiou et al. (2025). When contrasted against a non-semantic numerical task, semantic conditions caused either weak AG activation or no difference relative to the non-semantic baseline. However, when compared against the resting baseline, substantial *negative deactivation* was found, indicating a reversal of polarity in AG activation caused by a change of comparison baseline. This pattern also indicates the inherent ambiguity of using non-semantic tasks as a baseline: without a reference point (e.g., rest), it is difficult to determine whether semantic effects come from true activation (semantic > rest) or attenuated deactivation (i.e., semantic being less negative than non-semantic). (**B**) Crucially, certain paradigms induce robust AG activation irrespective of baselines used. For example, the social semantic task in Chiou et al. (2020) elicited robust AG activation regardless of a minimally semantic perceptual baseline or a highly semantic rest baseline. (**C**) This figure illustrates AG engagement along a continuum from perceptually *coupled* to perceptually *decoupled* cognition. At one end, externally oriented tasks that emphasise sensory-motor processing typically induce AG deactivation; at the other hand, introspective paradigms involving rich semantic content and information buffering (e.g., autobiographical recall and social evaluation tasks) robustly activate the AG, often exceeding the resting baseline. Situated between these extremes are semantic decisions on externally presented stimuli, common in task-based neuroimaging, which yield more ambiguous patterns – sometimes exceeding rest but frequently manifesting as deactivation or null difference.

One possible explanation for these patterns of AG activity is that this area tracks behavioural effort, with lower task demands associated with heightened AG engagement. This account appears to provide a parsimonious explanation for the data of Chiou et al. (2025): RTs for concrete words (1565 ms) were faster than those in the numerical task (1821 ms; *t*_(24)_ = 8.5, *p* < 0.001), whereas RTs for abstract words (1775 ms) did not differ from numerical RTs (*t*_(24)_ = 1.6, *p* = 0.13). This was also observed in Hoffman et al. (2015): RTs for concrete words (1790 ms) were substantially faster than numerical RTs (2138 ms; *t*_(17)_ = 8.26, *p* < 0.001), while RTs for abstract words (2156 ms) did not differ from numerical RTs (*t*_(17)_ = 0.65, *p* = 0.46). The same pattern was also evident in the social-cognition task of Chiou et al. (2020): both the Self and Queen conditions elicited faster RTs (Self: 1071 ms; Queen: 1056 ms) relative to the non-semantic perceptual task (1392 ms; both *p*s < 0.001). Across all three studies, the behaviourally easier conditions elicited stronger AG activity relative to those more effortful ones. However, the relationship between cognitive effort and AG activity is clearly more nuanced than a simplistic “low effort = high activation” account. A growing body of evidence has demonstrated that AG activity can substantially increase during demanding situations, provided that these tasks rely heavily on memory-guided cognition. For example, AG activation has been found to be substantially greater during short-term memory retention (Murphy *et al*., 2018; Murphy *et al*., 2019), episodic recall (Humphreys *et al*., 2022), and autobiographical recollection (Chiou *et al*., 2020), even though these tasks produced substantially slower behavioural RTs than perception-oriented control tasks. Moreover, in Chiou et al. (2020), both the Self and Queen tasks elicited greater AG activity than the behaviourally “effortless” resting baseline, further arguing against a simplistic interpretation in which AG engagement merely reflects reduced cognitive effort. Instead, these findings suggest that AG activity is jointly shaped by both cognitive effort and the extent to which cognition is memory-dependent and perceptually decoupled.

These results paint a clear picture: because baseline contexts differ in semantic richness, the choice of baseline can reverse the polarity of AG activation and thereby alter its interpretation; however, certain tasks reliably surpass varying baselines (even when they are behaviourally harder), revealing the types of processing that most potently engage the AG. As illustrated in Fig. 7C, task paradigms can be organised along a continuum of AG engagement. At one end, externally oriented tasks that emphasise sensory-motoric input/output and place minimal demands on semantic memory (e.g., perception or arithmetic) typically induce robust AG deactivation. At an intermediate level, semantic decision on externally presented stimuli (e.g., semantic similarity judgments that we used) engages the AG more than non-semantic tasks, yet often fail to surpass the rest baseline, as the need to process external stimuli and generate responses may dampen/disrupt ongoing semantic thoughts. At the opposite end, paradigms that rely heavily on introspectively directed cognition and afford extended periods for disengagement from external action/perception (e.g., autobiographical recall, narrative comprehension, and personality assessment) drive strong AG activation, often exceeding the semantically rich rest. Together, these patterns suggest that AG engagement scales with the degree to which cognition is internally guided, representationally rich, and perceptually decoupled.

## Discussion

The present investigation addresses a long-standing debate about how the AG contributes to semantic cognition: does its transmodal topography predispose the AG toward abstract semantics, or does greater AG activity for concrete words truly reflect concreteness? To this end, we conducted a multimethod investigation to identify the neurocognitive factors shaping AG function. In Study 1, cTBS to the AG disproportionately impaired abstract semantics and short-term mnemonic retention, suggesting that abstract semantics relies more heavily on AG resources. Furthermore, AG disruption during encoding also impaired subsequent memory retrieval. In Study 2, although concrete words initially elicited greater AG activity, this effect disappeared after controlling for RT and accuracy, indicating that the concreteness-bias may instead reflect cognitively effortless states. In Study 3, despite deactivation for abstract words, the AG sustained connectivity with “semantic control” areas (left IFG and pMTG), suggesting continued integration with the semantic network rather than disengagement. In Study 4, MDES revealed that concrete words were linked to experiences of imagery and automaticity, providing phenomenological accounts of elevated AG engagement for concrete words. Finally, Study 5 showed that baseline choices can flip the polarity of AG activation, while also demonstrating that AG recruitment is strongest during memory-guided cognition.

Taken together, our findings identify a set of interacting dimensions that offer a principled account of AG function and help resolve an ongoing debate on its role in semantic cognition. Rather than preferentially encoding concrete semantics, the AG contributes to cognition through mechanisms related to abstraction, information buffering, automaticity, and imagery. These dimensions tend to co-occur under cognitively less demanding contexts, giving rise to apparent concreteness effects that reflect accompanying mental states rather than concreteness *per se*. Critically, this framework clarifies both how the AG contributes to semantic cognition and how heightened AG activity for concrete words should be interpreted. We consider the implications of our results below.

### Representational abstractness in the default mode network

A large body of work (for review, Huntenburg *et al*., 2018; Jung *et al*., 2022) indicates that the default mode network generally (and the AG specifically) is preferentially engaged by forms of cognitive activities that are relatively immune to sensory-level changes, which suggests sensitivity to crossmodal semantic content that transcends superficial perceptual features. The AG is one of the regions found to exhibit such crossmodal sensitivity (Kuhnke *et al*., 2020a; Fernandino *et al*., 2022) although how its functionality differs from other crossmodal regions, such as the ATL (Lambon Ralph *et al*., 2017), remains contentious. From a connectomic perspective, this bias may reflect the AG’s position along connectomic gradients (Margulies *et al*., 2016). Topographically, the AG lies near the transmodal apex of the principal gradient, far from primary sensory cortices. This distance from sensory cortices reduces sensory/motoric constraint on its processing (Wang *et al*., 2024) and provides a neural estate where representations become liberated from sensory influences (Seghier, 2013). Under this framework, abstract semantics relies on cortical sections on the transmodal end of the gradient, while concrete semantics still retain stronger links to embodied representations in sensory regions (Meteyard *et al*., 2012). Consequently, abstract semantics depend more heavily on the integrity of transmodal zones (such as the AG), whereas concrete semantics can additionally draw on modality-specific systems and hence become comparatively less affected when the AG is disrupted by cTBS. This distinction offers a parsimonious account of the TMS findings in Study 1: Although AG disruption impaired both types of semantic processing, abstract semantics was more vulnerable because it relied more heavily on AG-mediated operations on semantic information when participants assessed abstract concepts. Importantly, while our discussion situates the AG near the apex of cortical hierarchy and emphasises its affiliation with the default mode network, it is important to recognise that the AG comprises multiple subregions within the inferior parietal lobule and lies in proximity to the dorsal/ventral attention and frontoparietal networks (Yeo *et al*., 2011), with distinct subregions showing partially different patterns of large-scale network connectivity. Converging evidence indicates that different inferior parietal subregions show distinct patterns of coupling with functional systems: Anterior portions (supramarginal gyrus) more closely linked to the ventral attention and frontoparietal networks; dorsal portions (intraparietal sulcus) linked to dorsal attention network; posterior portions (AG) more associated with the default mode network (Numssen *et al*., 2021; Williams *et al*., 2022). Such heterogeneity is consistent with the view that AG function emerges from dynamic interactions across multiple large-scale networks rather than from a unitary network affiliation. Embedded within this heterogenous structure-function mapping, the present findings suggest that abstract semantic processing may place greater demands on the neural resources of middle AG owing to its intermediate locus that spans semantic/language and attention/control systems.

### Information buffering in the AG during temporally evolving cognition

Semantic decision on single-word stimuli has been shown to engage the AG (e.g., Kuhnke *et al*., 2023b), although the overall pattern of evidence remains equivocal (Humphreys *et al*., 2015). Compared to the divergent results in the single-word literature, there is overwhelming evidence that temporally evolving cognitive activities, such as reading comprehension, sentence composition, episodic/autobiographic memory recollection, robustly engage the AG (e.g., Boylan *et al*., 2015; Branzi *et al*., 2020; Humphreys *et al*., 2024; Demirkan and Branzi, 2025). Notably, AG recruitment is observed not only during temporally extended cognitive activities, but also during simple short-term maintenance, such as recalling a stimulus presented moments earlier in a one-back task (Murphy *et al*., 2018; Murphy *et al*., 2019), as well as during contexts wherein semantic messages need to be integrated across time to form a combinatorial meaning (Price *et al*., 2015; Branzi *et al*., 2020; Lanzoni *et al*., 2020). Building on extensive evidence regarding AG involvement in temporally extended cognition, the AG has been proposed to support a domain-general buffering mechanism that transiently maintains online information as it unfolds over time (Humphreys *et al*., 2021; Humphreys and Tibon, 2023). On this account, the AG does not store semantic or episodic content *per se* but temporarily retains task-relevant representations to support ongoingly evolving operation during a task, such as constructing sentence-level meaning across words. This buffering enables the integration of information over successive inputs, allowing short-term memory buffer to be constantly updated. Consequently, the AG is more reliably engaged by cognitive activities that require the accrual and maintenance of information over time. Consistent with this view, our finding that cTBS to the AG disproportionately disrupted contexts with buffering requirement provides causal support for the AG’s role in memory-guided cognition. Together with our evidence of a greater disruption on abstract semantics, we show that both buffering and abstraction engage a shared AG computation that keeps information in *representationally abstract* and *operationally buffered* formats, relative to perceptually anchored representations tied to sensory-motoric systems.

### Automaticity and mental imagery

Another critical feature of AG is its sensitivity to task automaticity. AG engagement scales with familiar stimuli/rules that enable automated reaction/retrieval than effortful executive control. (Davey *et al*., 2015; Shamloo and Helie, 2016; Vatansever *et al*., 2017). Such a propensity towards automaticity suggests that the AG, as part of the default mode network, is heavily involved in memory-guided cognition by leveraging prior knowledge and familiarity, such that its engagement is greatest when existing memory (be it explicit semantic knowledge or implicit procedural schema) can be applied with minimal need for executive control. Given that concrete words are acquired much earlier in life than abstract words and are therefore more familiar (Della Rosa *et al*., 2010), this account provides a principled explanation for their effort-light processing, a pattern that is mirrored in our MDES data showing predominant experiences of cognitive ease for concrete words and heightened effort for abstract words. Furthermore, our MDES findings revealed that participants engaged in significantly more mental simulation when processing concrete words, likely because concrete concepts more readily evoke vivid mental images (Altarriba *et al*., 1999), consistent with their well-documented imageability advantage and with reliable AG engagement during mental imagery (e.g., Boccia *et al*., 2015; Spreng *et al*., 2018).

### Methodological insights and future directions

Our results offer a key methodological insight that sharpens interpretation of AG responses. Baseline choice profoundly shapes apparent activation: positive ‘semantic > non-semantic’ effects are not inherently diagnostic, as they may reflect either genuine recruitment or reduced deactivation relative to externally oriented control tasks. Introducing rest as a cautiously interpreted reference point helps resolve this ambiguity and identifies paradigms that truly exceed the resting baseline. Consistent with the perceptual decoupling account of Jefferies and Smallwood (2025), robust AG engagement emerges when cognition is internally guided, conceptually rich, and shielded from sensory interruption – conditions that support sustained buffering and immersive semantic integration beyond immediate perceptual demands. In addition to baseline selection, there are other methodological and theoretical issues worth noting. First, the AG is a heterogeneous brain structure. As mentioned earlier, the AG comprises multiple subregions with distinct connectomic fingerprints and functional profiles (Caspers and Zilles, 2018; Numssen *et al*., 2021; Humphreys *et al*., 2022). To ensure consistency and direct comparability across studies, in the present study we therefore adopted a single coordinate for AG localisation, allowing fMRI analyses to be aligned with the cTBS target. However, when cross-methods comparability is not required, finer-grained parcellation of parietal subregions is certainly preferable, and data-driven approach (such as k-means clustering) would especially be suited to investigate the fractionation within the AG (e.g., Numssen *et al*., 2021). Moreover, adopting fMRI-guided TMS combined with individual electric-field modelling (Kuhnke *et al*., 2025), potentially with concurrent MDES, may provide valuable insight about how TMS modulates ongoing thought patterns. Second, while this study focuses on univariate analyses, multivariate decoding can detect representational pattern subtlety that univariate methods may miss. For example, decoding has been used to show that AG neural patterns encode information about concrete (Fernandino *et al*., 2022), abstract (Kaiser *et al*., 2022), and person-related semantics (Chiou *et al*., 2023). Note that, decoding does not indicate the direction of effects (i.e., the AG favours one task over another); instead, it shows AG neural patterns are distinguishable by an algorithm. If interpreted carefully, multivariate approaches may provide a complementary window into the informational content of the AG beyond activation magnitude. Finally, the division of labour between the AG and ATL is an important topic for future research. It has been well-established that the ATL is one of the key neural substrates for encoding semantic knowledge (e.g., Chiou and Ralph, 2018; Chiou and Lambon Ralph, 2019; Cox *et al*., 2024), while the AG has also been implicated in many semantic tasks (for review, Kuhnke *et al*., 2023a). While beyond the scope of the present study, clarifying how the AG and ATL differentially and complementarily contribute to semantic cognition will be crucial for understanding how semantic representations are encoded, integrated, buffered, and flexibly deployed across transmodal zones of the cortex.

## Acknowledgement

**Study 1** was supported by a British Academy Small Research Grant to the author RC (SRG2324\240157). **Study 2**, originally reported in Chiou et al. (2025), was supported by an MRC Programme Grant and Intramural Funding to Matthew A. Lambon Ralph (MR/R023883/1; MC_UU_00005/18), together with a Sir Henry Wellcome Fellowship to RC (201381/Z/16/Z). **Study 3**, originally reported in Hoffman et al. (2015), was supported by an MRC Programme Grant to MALR (MR/J004146/1). **Study 4** was supported by a British Academy Small Research Grant to RC (SRG2324\240157). **Study 5**, originally reported in Chiou et al. (2020), was supported by an MRC Programme Grant to MALR (MR/R023883/1) and a Sir Henry Wellcome Fellowship to RC (201381/Z/16/Z). We thank Matthew Lambon Ralph, Paul Hoffman, and Richard Binney for sharing the fMRI dataset that enabled the re-analyses reported here. We are grateful to Matthew Lambon Ralph for his comments on an earlier version of the manuscript. We also thank Olivier Pacaud, Sagarika Saproo, Seemeen Bokth, Bronwyn Woods, Tania Nazarian, and Rahim Ahmed for assisting with data collection, laboratory setup, and participant recruitment of the neurostimulation and experiential sampling experiments. Finally, we thank the two anonymous reviewers for their helpful suggestions and for raising issues that we had not considered.

