## Supplemental information for "Resolving the Abstract–Concrete Paradox in the Angular Gyrus: A Multimethod Investigation"

**Table of contents**:

Supplemental Table 1 – The reaction times and accuracy rates of all conditions in Study 1 (Page 2)
Supplemental Table 2 – Linear mixed effects on the TMS data Study 1: LME codes (Page 3 – 13)
Supplemental Table 3 – Linear mixed effects on the TMS data Study 1: LME results (Page 14 – 15)
Supplemental Table 4 – Linear mixed effects and Bayes testing on the fMRI data Study 2: LME codes (Page 16)
Supplemental Table 5 – Psychophysiological interaction analyses on Study 3 (Page 17)
Supplemental Figure 1 – The scree plot of principal component analysis in Study 4 (Page 18)
Supplemental Figure 2 – Percentage signal change figures of Study 5 (Page 19)

### Supplemental Table 1

Reaction time data (millisecond) of all conditions of the continuous theta-burst stimulation experiment in Study 1

| | Angular gyrus | | | | | | | | | | | |
|---|---|---|---|---|---|---|---|---|---|---|---|---|
| | Concrete | | | | | | Abstract | | | | | |
| | Related | | | Unrelated | | | Related | | | Unrelated | | |
| | SimVis | VisVis | SpchVis | SimVis | VisVis | SpchVis | SimVis | VisVis | SpchVis | SimVis | VisVis | SpchVis |
| *Mean* | 1173 | 937 | 934 | 1396 | 1038 | 1011 | 1448 | 1205 | 1174 | 1721 | 1304 | 1216 |
| *SEM* | 24 | 22 | 21 | 34 | 21 | 28 | 42 | 28 | 34 | 38 | 44 | 37 |

| | Vertex | | | | | | | | | | | |
|---|---|---|---|---|---|---|---|---|---|---|---|---|
| | Concrete | | | | | | Abstract | | | | | |
| | Related | | | Unrelated | | | Related | | | Unrelated | | |
| | SimVis | VisVis | SpchVis | SimVis | VisVis | SpchVis | SimVis | VisVis | SpchVis | SimVis | VisVis | SpchVis |
| *Mean* | 1138 | 869 | 852 | 1329 | 929 | 904 | 1386 | 1058 | 1016 | 1584 | 1163 | 1080 |
| *SEM* | 33 | 26 | 25 | 35 | 22 | 25 | 33 | 25 | 27 | 38 | 30 | 22 |

Accuracy rate data (% correct) of all conditions of the continuous theta-burst stimulation experiment in Study 1

| | Angular gyrus | | | | | | | | | | | |
|---|---|---|---|---|---|---|---|---|---|---|---|---|
| | Concrete | | | | | | Abstract | | | | | |
| | Related | | | Unrelated | | | Related | | | Unrelated | | |
| | SimVis | VisVis | SpchVis | SimVis | VisVis | SpchVis | SimVis | VisVis | SpchVis | SimVis | VisVis | SpchVis |
| *Mean* | 95 | 94 | 93 | 97 | 97 | 98 | 86 | 78 | 82 | 93 | 95 | 95 |
| *SEM* | 1.19 | 0.68 | 1.16 | 0.96 | 1.49 | 0.81 | 1.77 | 1.78 | 2.04 | 1.94 | 1.29 | 1.09 |

| | Vertex | | | | | | | | | | | |
|---|---|---|---|---|---|---|---|---|---|---|---|---|
| | Concrete | | | | | | Abstract | | | | | |
| | Related | | | Unrelated | | | Related | | | Unrelated | | |
| | SimVis | VisVis | SpchVis | SimVis | VisVis | SpchVis | SimVis | VisVis | SpchVis | SimVis | VisVis | SpchVis |
| *Mean* | 95 | 94 | 92 | 97 | 98 | 97 | 84 | 82 | 82 | 92 | 96 | 96 |
| *SEM* | 0.82 | 1.23 | 0.93 | 0.77 | 0.99 | 0.72 | 1.84 | 2.41 | 2.30 | 1.57 | 1.16 | 1.09 |

### Supplemental Table 2 – The codes of linear-mixed effects modelling on the TMS data of Study 1

```r
# ========================================================================
# Codes on this page are just for loading file and setting up parameters
# ========================================================================

# 1. LOAD LIBRARIES
library(lme4)
library(lmerTest)
library(readxl)
library(emmeans)
library(dplyr)

# 2. IMPORT DATA
my_data <- read_excel("TMS_data_formatted_for_LME - (Session added).xlsx")

# 3. PREPARE VARIABLES
my_data <- my_data %>%
  mutate(
    Subject = factor(Subject),
    TMS = factor(TMS, levels = c("Vertex", "AG")),
    Concreteness = factor(Concreteness),
    Relatedness = factor(Relatedness),
    Presentation = factor(Presentation),
    Session = factor(Session, levels = c("First", "Second"))
  )

# Check coding
str(my_data)
table(my_data$Subject, my_data$TMS, my_data$Session)

# 4. SET OPTIMISER
ctrl <- lmerControl(
  optimizer = "bobyqa",
  optCtrl = list(maxfun = 100000)
)
```

```r
# ===========================================================================
# PART A: TESTING IF 'SESSION' (FIRST or SECOND) HAS ANY INFLUENCE ON RESULTS
# ===========================================================================

# This model does not contain Session as a fixed effect.
model_no_session <- lmer(
  RT ~ TMS * Concreteness * Relatedness * Presentation +
    (1 | Subject),
  data = my_data,
  REML = FALSE,
  control = ctrl
)

# This model includes Session and tests its main effect.
model_session_main <- lmer(
  RT ~ TMS * Concreteness * Relatedness * Presentation +
    Session +
    (1 | Subject),
  data = my_data,
  REML = FALSE,
  control = ctrl
)

# This model tests the Session main effect & Session x TMS interaction.
model_session_by_TMS <- lmer(
  RT ~ TMS * Concreteness * Relatedness * Presentation +
    Session * TMS +
    (1 | Subject),
  data = my_data,
  REML = FALSE,
  control = ctrl
)
```

```r
# This model includes all possible interactions between Session and other factors.
model_session_all_interactions <- lmer(
  RT ~ TMS * Concreteness * Relatedness * Presentation +
    Session * (TMS + Concreteness + Relatedness + Presentation) +
    (1 | Subject),
  data = my_data,
  REML = FALSE,
  control = ctrl
)

# Compare whether Session improves the model
fixed_model_comparison <- anova(
  model_no_session,
  model_session_main,
  model_session_by_TMS,
  model_session_all_interactions
)

print(fixed_model_comparison)
print(AIC(model_no_session, model_session_main, model_session_by_TMS, model_session_all_interactions))
print(BIC(model_no_session, model_session_main, model_session_by_TMS, model_session_all_interactions))
```

```r
# ============================================================================
# PART B: EXPLORING DIFFERENT STRUCTURES OF RANDOM EFFECTS & MODEL SELECTION
# ============================================================================

# Following the reviewer's recommendation, we evaluated a series of increasingly complex
# random-effect structures to account for inter-individual variability in task performance.
# We compared models with subject-specific random slopes for TMS, Concreteness, Session, Relatedness, & Presentation.
# Both correlated (|) and uncorrelated (||) structures of random effects were considered. The uncorrelated models were
# included simply as alternative models in cases where more complex correlated random-effect structures might result in
# singular fits or convergence difficulties. Correlated models allowed correlations among task-related random effects to be
# estimated freely, whereas uncorrelated models included the same random effects but constrained their correlations to zero.

# Below we first specify 5 models with different random-effect structures

# Simplest model with only random intercepts for subjects.
m0 <- lmer(
  RT ~ TMS * Concreteness * Relatedness * Presentation +
    Session +
    (1 | Subject),
  data = my_data,
  REML = FALSE,
  control = ctrl
)

# Slightly more complex model with random slopes for subjects and TMS.
m1_TMS <- lmer(
  RT ~ TMS * Concreteness * Relatedness * Presentation +
    Session +
    (1 + TMS | Subject),
  data = my_data,
  REML = FALSE,
  control = ctrl
)
```

```r
# Even more complex model with random slopes for subjects, TMS, and Session.
m2_TMS_Session <- lmer(
  RT ~ TMS * Concreteness * Relatedness * Presentation +
    Session +
    (1 + TMS + Session | Subject),
  data = my_data,
  REML = FALSE,
  control = ctrl
)

# Further complex model with random slopes for subjects, TMS, Session, and Relatedness.
m3_TMS_Session_Relatedness <- lmer(
  RT ~ TMS * Concreteness * Relatedness * Presentation +
    Session +
    (1 + TMS + Session + Relatedness | Subject),
  data = my_data,
  REML = FALSE,
  control = ctrl
)

# Most complex model with random slopes for subjects, TMS, Concreteness, Session, Relatedness, and Presentation.
m4_TMS_Concreteness_Session_Relatedness_Presentation <- lmer(
  RT ~ TMS * Concreteness * Relatedness * Presentation +
    Session +
    (1 + TMS + Concreteness + Session + Relatedness + Presentation | Subject),
  data = my_data,
  REML = FALSE,
  control = ctrl
)
```

```r
# Below are uncorrelated versions of the random-effect models (M1 – M4).

m2_uncorrelated <- lmer(
  RT ~ TMS * Concreteness * Relatedness * Presentation +
    Session +
    (1 + TMS + Session || Subject),
  data = my_data,
  REML = FALSE,
  control = ctrl
)

m3_uncorrelated <- lmer(
  RT ~ TMS * Concreteness * Relatedness * Presentation +
    Session +
    (1 + TMS + Session + Relatedness || Subject),
  data = my_data,
  REML = FALSE,
  control = ctrl
)

m4_uncorrelated <- lmer(
  RT ~ TMS * Concreteness * Relatedness * Presentation +
    Session +
    (1 + TMS + Concreteness + Session + Relatedness + Presentation || Subject),
  data = my_data,
  REML = FALSE,
  control = ctrl
)
```

```
# Check convergence and singularity of the eight models as necessary steps

model_names <- c(
  "m0",                                                   # m0: random slope for Subj
  "m1_TMS",                                               # m1: random slope for Subj & TMS
  "m2_TMS_Session",                                       # m2: random slope for Subj, TMS, & Sess
  "m3_TMS_Session_Relatedness",                           # m3: random slope for Subj, TMS, Sess, & Rel
  "m4_TMS_Concreteness_Session_Relatedness_Presentation", # m4: random slope for Subj, TMS, Conc, Sess, Rel, & Pres
  "m2_uncorrelated",                                      # m5: similar to m2 but it is an uncorrelated version of m2
  "m3_uncorrelated",                                      # m6: similar to m3 but it is an uncorrelated version of m3
  "m4_uncorrelated"                                       # m7: similar to m4 but it is an uncorrelated version of m4
)

models <- list(
  m0,
  m1_TMS,
  m2_TMS_Session,
  m3_TMS_Session_Relatedness,
  m4_TMS_Concreteness_Session_Relatedness_Presentation,
  m2_uncorrelated,
  m3_uncorrelated,
  m4_uncorrelated
)

singularity_check <- data.frame(
  Model = model_names,
  Singular = sapply(models, isSingular)
)
```

```r
# Calculate and compare AIC and BIC for the eight different LME models

random_model_comparison_AIC <- AIC(
  m0,
  m1_TMS,
  m2_TMS_Session,
  m3_TMS_Session_Relatedness,
  m4_TMS_Concreteness_Session_Relatedness_Presentation,
  m2_uncorrelated,
  m3_uncorrelated,
  m4_uncorrelated
)

random_model_comparison_BIC <- BIC(
  m0,
  m1_TMS,
  m2_TMS_Session,
  m3_TMS_Session_Relatedness,
  m4_TMS_Concreteness_Session_Relatedness_Presentation,
  m2_uncorrelated,
  m3_uncorrelated,
  m4_uncorrelated
)

print(random_model_comparison_AIC)
print(random_model_comparison_BIC)

# Likelihood-ratio tests for nested models
anova(m0, m1_TMS, m2_TMS_Session, m3_TMS_Session_Relatedness, m4_TMS_Concreteness_Session_Relatedness_Presentation)
anova(m0, m2_uncorrelated, m3_uncorrelated, m4_uncorrelated)
```

```r
# Select the final winner model based on the smallest AIC/BIC values

# Rank models by AIC - just rank them from smallest to largest
random_model_comparison_AIC[order(random_model_comparison_AIC$AIC), ]

# Rank models by BIC - just rank them from smallest to largest
random_model_comparison_BIC[order(random_model_comparison_BIC$BIC), ]


# SAVE OUTPUTS

write.csv(as.data.frame(random_model_comparison_AIC),
          "Random_Effects_Model_Comparison_AIC.csv")

write.csv(as.data.frame(random_model_comparison_BIC),
          "Random_Effects_Model_Comparison_BIC.csv")
```

```r
# ==========================================================================================
# PART C: RUNNING LME ANALYSIS TO EXAMINE THE MAIN/INTERACTION EFFECTS OF THE WINNING MODEL
# ==========================================================================================

# Based on model section, we found that Model 4, where we included random intercepts for participant-level variance,
# as well as random slopes for all experimental variables, was found to have consistently smaller AIC and BIC values.
# Therefore, we conducted LME to examine the main/interaction effects and post-hoc comparisons for this winner model.

# 1. LOAD LIBRARIES (already installed)
library(lme4)
library(lmerTest)
library(readxl)
library(emmeans) # Adding this for post-hoc tests

# 2. IMPORT DATA
# Ensure "experiment_data_long_format.xlsx" is in your working directory
my_data <- read_excel("TMS_data_long_format_for_LME - (Session added).xlsx")

# 3. PREPARE FACTORS
# Converting text columns into "Factors" so R treats them as experimental groups
my_data$Subject      <- as.factor(my_data$Subject)
my_data$TMS          <- as.factor(my_data$TMS)
my_data$Session      <- as.factor(my_data$Session)
my_data$Concreteness <- as.factor(my_data$Concreteness)
my_data$Relatedness  <- as.factor(my_data$Relatedness)
my_data$Presentation <- as.factor(my_data$Presentation)

# Check coding
str(my_data); table(my_data$Subject, my_data$TMS, my_data$Session)

# Set optimizer
ctrl <- lmerControl(optimizer = "bobyqa", optCtrl = list(maxfun = 1e5))
```

```r
# 3. RUN THE LINEAR MIXED MODEL
m4_TMS_Concreteness_Session_Relatedness_Presentation <- lmer(
  RT ~ TMS * Concreteness * Relatedness * Presentation +
    Session +
    (1 + TMS + Concreteness + Session + Relatedness + Presentation | Subject),
  data = my_data,
  REML = FALSE,
  control = ctrl
)

# 4. VIEW MAIN RESULTS (ANOVA Table)
anova_results <- anova(m4_TMS_Concreteness_Session_Relatedness_Presentation)
print(anova_results)

# 5. POST-HOC TESTS
posthoc_concreteness <- emmeans(m4_TMS_Concreteness_Session_Relatedness_Presentation, pairwise ~ TMS | Concreteness)
posthoc_presentation <- emmeans(m4_TMS_Concreteness_Session_Relatedness_Presentation, pairwise ~ TMS | Presentation)
print(posthoc_concreteness)
print(posthoc_presentation)

# 6. SAVE RESULTS TO EXCEL-FRIENDLY FILE
write.csv(as.data.frame(anova_results), "Final_Results_of_the_Winner.csv")
```

### Supplemental Table 3 – Results of linear-mixed effects modelling on the TMS data of Study 1

#### (1) Part A – including '*Session*' as a fixed-effect variable

| Models | npar | AIC | BIC | logLik | -2logLik | Chisq | Df | Pr(>Chisq) |
|---|---|---|---|---|---|---|---|---|
| Model_no_session | 26 | 7546.082 | 7659.340 | -3747.041 | 7494.082 | - | - | - |
| Model_session_main | 27 | 7548.036 | 7665.651 | -3747.018 | 7494.036 | 0.045640301 | 1 | 0.830830995 |
| Model_session_by_TMS | 28 | 7549.399 | 7671.370 | -3746.699 | 7493.399 | 0.637107064 | 1 | 0.424760317 |
| Model_session_all_interactions | 32 | 7554.492 | 7693.887 | -3745.246 | 7490.492 | 2.90678177 | 4 | 0.573544482 |

Both AIC and BIC favoured the model without Session, indicating that adding Session-related terms did not improve model fit. Consistent with this, likelihood-ratio Chi-square tests showed that none of the models incorporating Session significantly outperformed the baseline model without Session. The sections relevant to statistical interpretation are highlighted with pink.

#### (2) Part B – investigating different structures of random effects

| Models | Df | AIC | BIC |
|---|---|---|---|
| Model 0 - Participants as the only random effect | 27 | 7548.036 | 7665.651 |
| Model 1 - Participant and TMS as random effects | 29 | 7486.921 | 7613.248 |
| Model 2 - Participant, TMS, Session as random effects | 32 | 7484.109 | 7623.504 |
| Model 3 - Participant, TMS, Session, Relatedness as random effects | 36 | 7452.525 | 7609.345 |
| Model 4 - Participant, TMS, Concreteness, Session, Relatedness, Presentation as random effects | 47 | 7357.430 | 7592.660 |
| Model 5 - Very similar to Model 2 but this model assumes *no correlation* between factors | 33 | 7493.943 | 7637.695 |
| Model 6 - Very similar to Model 3 but this model assumes *no correlation* between factors | 36 | 7463.581 | 7620.401 |
| Model 7 - Very similar to Model 4 but this model assumes *no correlation* between factors | 42 | 7363.059 | 7559.084 |

Comparison of alternative random-effect structures showed a general improvement in model fit as additional subject-specific random slopes were included in the model, reflected by progressively lower AIC and BIC values. This finding supports modelling inter-individual variability in responses to the experimental factors. Model 4 and 7 showed overall smaller AIC and BIC values (sections highlighted with pink) as compared to other models. Particularly, Model 4 had the smallest AIC value amongst all models. Thus, we concluded that Model 4 was the winner from this comparison process.

#### (3) Part C – examining the LME results of the winner (Model 4)

| Different effects | Sum Sq | Mean Sq | NumDF | DenDF | F value | Pr(>F) |
|---|---|---|---|---|---|---|
| *TMS* | *190869.30* | *190869.30* | *1* | *10.79* | *17.00* | *0.002* |
| Concreteness | 2332020.19 | 2332020.19 | 1 | 24.01 | 207.65 | 0.000 |
| Relatedness | 396523.88 | 396523.88 | 1 | 24.02 | 35.31 | 0.000 |
| Presentation | 2164640.43 | 1082320.22 | 2 | 27.98 | 96.37 | 0.000 |
| Session | 4703.93 | 4703.93 | 1 | 16.28 | 0.42 | 0.527 |
| *TMS:Concreteness* | *99172.47* | *99172.47* | *1* | *446.13* | *8.83* | *0.003* |
| TMS:Relatedness | 20330.03 | 20330.03 | 1 | 446.13 | 1.81 | 0.179 |
| Concreteness:Relatedness | 5877.79 | 5877.79 | 1 | 446.13 | 0.52 | 0.470 |
| *TMS:Presentation* | *61566.71* | *30783.36* | *2* | *446.13* | *2.74* | *0.066* |
| Concreteness:Presentation | 150542.88 | 75271.44 | 2 | 446.13 | 6.70 | 0.001 |
| Relatedness:Presentation | 712402.86 | 356201.43 | 2 | 446.13 | 31.72 | 0.000 |
| TMS:Concreteness:Relatedness | 2550.25 | 2550.25 | 1 | 446.13 | 0.23 | 0.634 |
| TMS:Concreteness:Presentation | 190.19 | 95.10 | 2 | 446.13 | 0.01 | 0.992 |
| TMS:Relatedness:Presentation | 17665.59 | 8832.79 | 2 | 446.13 | 0.79 | 0.456 |
| Concreteness:Relatedness:Presentation | 11403.46 | 5701.73 | 2 | 446.13 | 0.51 | 0.602 |
| TMS:Concreteness:Relatedness:Presentation | 16144.97 | 8072.48 | 2 | 446.13 | 0.72 | 0.488 |

Following model selection, LME was conducted for the best-fitting model (Model 4), which included random slopes for all experimental variables, as well as random intercepts for participants, and assumed the correlations between experimental factors to be non-zero. Significant effects ($p < .05$) are highlighted in pink, while effects involving TMS are presented in red italics. Post-hoc comparisons were conducted for significant interactions involving TMS effects (as well as strong trends for exploratory purposes). These data are highly consistent with the results of ANOVA (see main text).

Although the results are presented in an *F*-test format, the tests were conducted within the LME framework using the Satterthwaite approximation for degrees of freedom. These tests assess the significance and robustness of the fixed effects within the best-supported mixed-effects structure and should not be interpreted as a conventional repeated-measures ANOVA.

### Supplemental Table 4 – The codes of linear-mixed effects modelling on the fMRI data of Study 2

```
# =========================================================
# Linear mixed effects on Study 2 (Chiou et al., 2025)
# =========================================================

# 1. LOAD LIBRARIES
library(lme4)
library(lmerTest)

# 2. IMPORT DATA
df <- read.csv("Data_file.csv")

# 3. PREPARE VARIABLES
df$Participant <- as.factor(df$Subject)
df$Condition   <- as.factor(df$Condition)

# 4. MODEL 1: Condition only (Concrete vs. Abstract) without ReactionTime
Model1 <- lmer(BrainActivity ~ Condition + (1 | Participant), data = df)
summary(Model1)

# 5. MODEL 2: Condition (Concrete vs. Abstract) + ReactionTime + Interaction
Model2 <- lmer(BrainActivity ~ Condition * ReactionTime + (1 | Participant), data = df)
summary(Model2)

# 6. Derive Bayes factor for the concreteness effect under different models
library(BayesFactor)
bf_withoutRT <- ttest.tstat(-4.964, 25)  # Under Model1 (without ReactionTime), t=-4.964, N=25
print(exp(bf_withoutRT$bf))
bf_withRT    <- ttest.tstat(-0.914, 25)  # Under Model2 (with ReactionTime), t=-0.914, N=25
print(exp(bf_withRT$bf))
```

**Supplemental Table 5 – Psychophysiological interaction analyses on the fMRI data of Study 3**

Standard psychophysiological analysis with SPM12 (Ashburner et al., 2014)

| Brain region | Cluster size (# of voxels) | Local peak (*Z*-value) | X | Y | Z |
|---|---|---|---|---|---|
| Inferior frontal gyrus (left) | | 5.65 | -42 | 32 | -6 |
| Posterior middle temporal gyrus (left) | 1758 | 4.87 | -58 | -50 | 12 |
| Middle superior temporal gyrus (left) | | 4.66 | -52 | -24 | -2 |

$P < 0.005$ (voxelwise); $P < 0.05$ (cluster-corrected for familywise error)

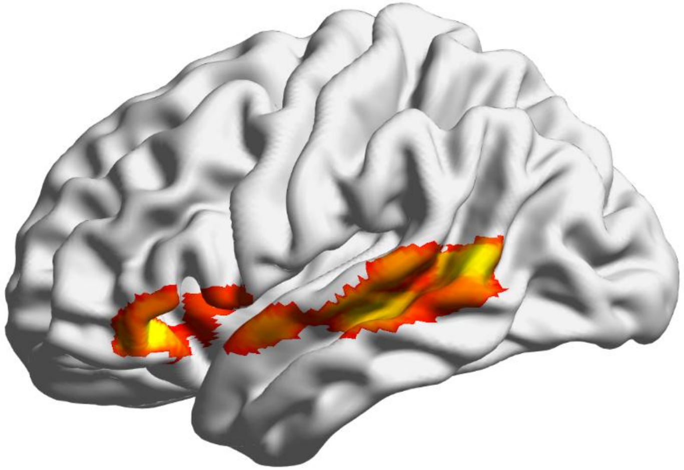

Generalised psychophysiological interaction analysis (gPPI; McLaren et al., 2012)

| Brain region | Cluster size (# of voxels) | Local peak (*Z*-value) | X | Y | Z |
|---|---|---|---|---|---|
| Posterior middle temporal gyrus (left) | | 5.23 | -58 | -52 | 10 |
| Anterior superior temporal gyrus (left) | 1559 | 4.74 | -58 | 2 | -6 |
| Inferior frontal gyrus (left) | | 4.25 | -50 | 46 | -4 |

$P < 0.005$ (voxelwise); $P < 0.05$ (cluster-corrected for familywise error)

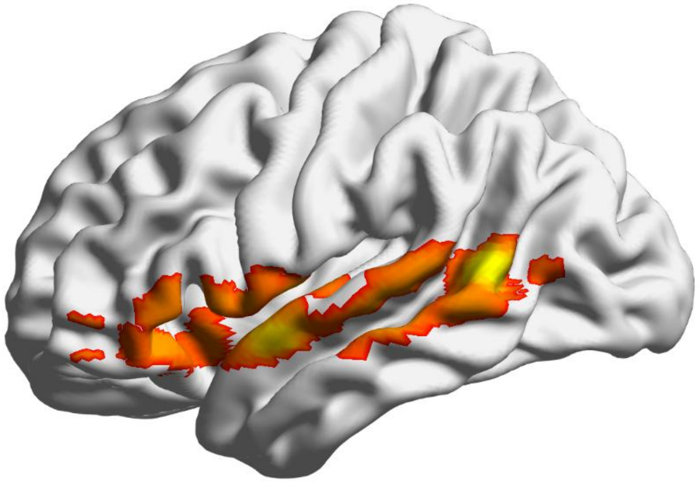

**Supplemental Figure 1 – Scree plot of the principal component analysis of Study 4**

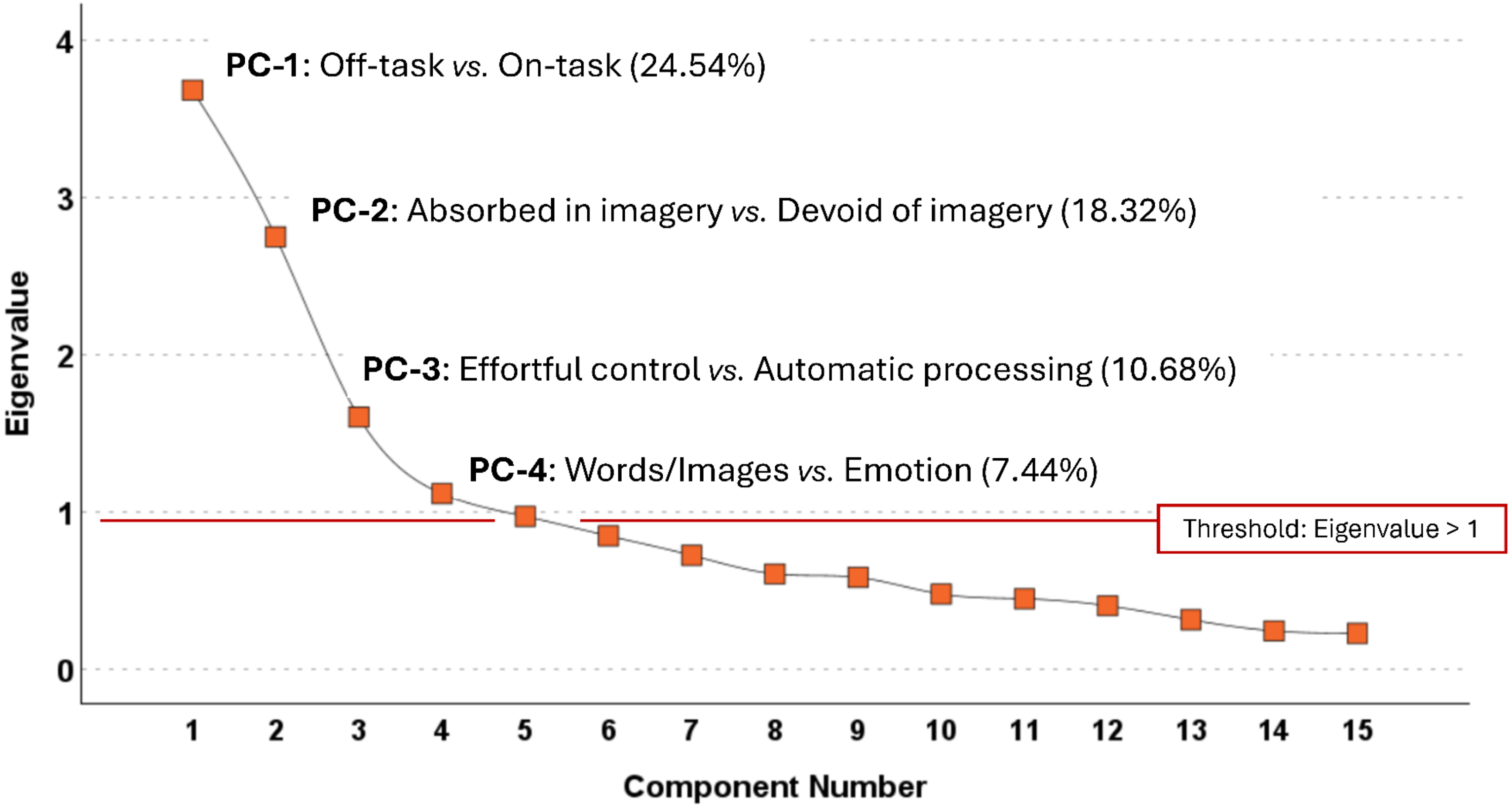


Using the default threshold of SPSS (eigenvalue equal to or greater than 1), followed by the tests of split-half reliability to ensure robustness of these components, the principal component analysis (PCA) identified four components surpassing the threshold. The values in the parentheses indicate the percentage of unique variance explained by each of the four significant components.

### Supplemental Figure 2 – Percentage signal change figures of Study 5

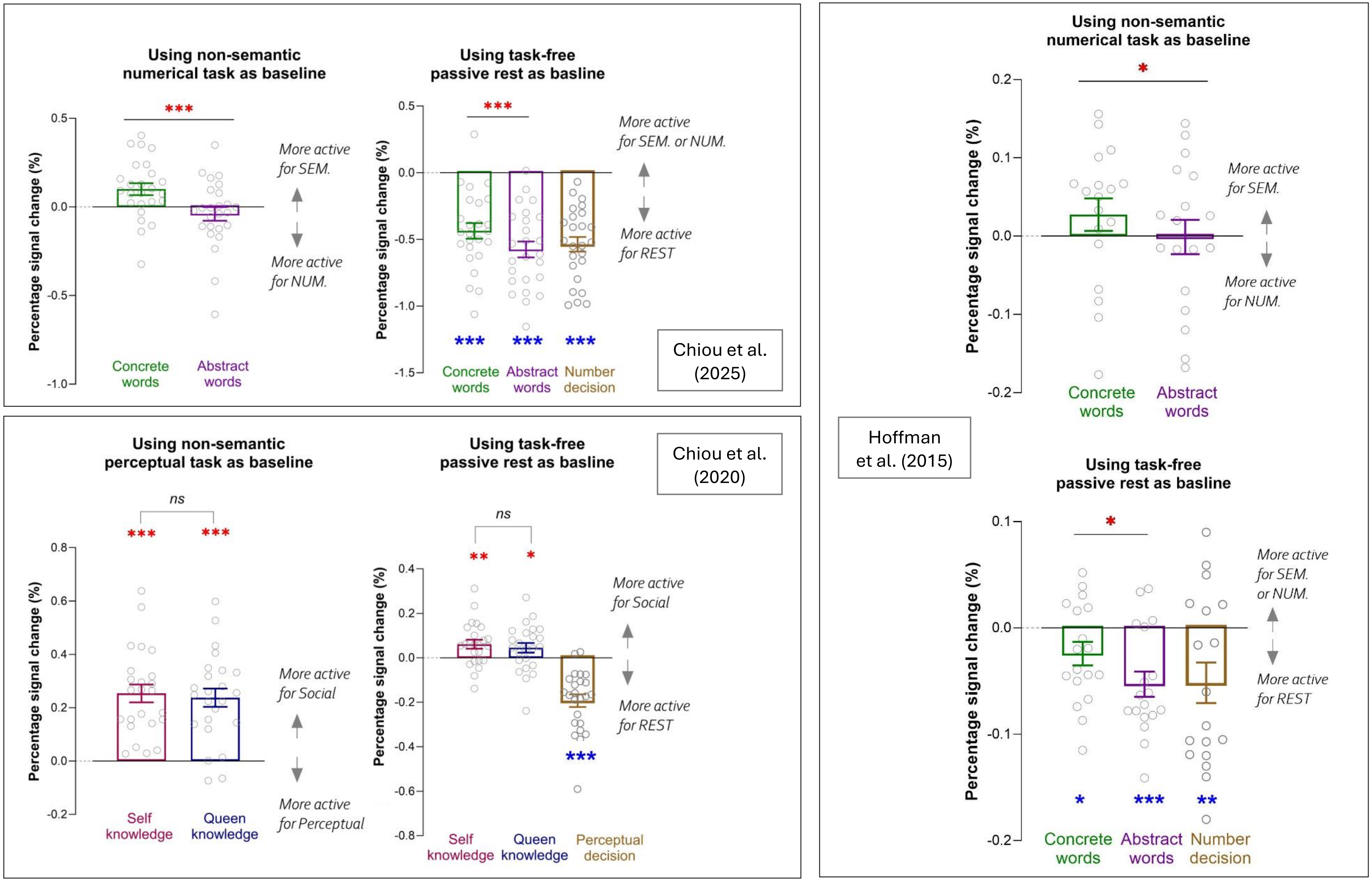
